# Protein-structure-sensitive mid-infrared optoacoustic microscopy enables label-free assessment of drug therapy in myeloma cells

**DOI:** 10.1101/2024.02.21.581391

**Authors:** Francesca Gasparin, Marlene R. Tietje, Eslam Katab, Aizada Nurdinova, Tao Yuan, Andriy Chmyrov, Nasire Uluç, Dominik Jüstel, Florian Bassermann, Vasilis Ntziachristos, Miguel A. Pleitez

## Abstract

Conventional live-cell optical microscopy lacks sensitivity and specificity for label-free detection of intracellular protein-structure dynamics, such as conformational transition from α-helix to β-sheet. Detecting intermolecular β-sheet formation, for instance, is important because it is a hallmark of misfolded proteins and aggresome formation—which are intrinsic indicators of cell apoptosis in myeloma therapy. Going beyond conventional optical microscopy, we introduce a single-cell imaging technology with label-free sensitivity to intracellular intermolecular β-sheet formation in living cells. This unique ability was attained by exploiting the spectral specificity of the mid-infrared amide I region (1700 – 1600 cm^-1^) to protein structure and the positive-contrast nature of optoacoustic microscopy. By means of this technology, we were able to monitor the efficiency of proteasome inhibition in a myeloma cell line and—as a first demonstration towards clinical translation—in biopsied myeloma cells from patients. Achieving label-free monitoring of treatment at a single-cell level allows longitudinal assessment of response heterogeneity, which could provide crucial therapeutic information, such as patient-specific sensitivity to treatment, thus facilitating personalized medicine in myeloma therapy.

## Introduction

The cellular ubiquitin-proteasome system (UPS) is responsible for the degradation of dysfunctional intracellular proteins, such as misfolded proteins resulting from α-helix to intermolecular β-sheet conformational transitions.^1^ When accumulated, misfolded proteins lead to cytotoxic aggresome formation that eventually triggers apoptosis.^1^ In cancer therapy, inhibition of the UPS to induce apoptosis is a common strategy for the treatment of, for example, multiple myeloma, amyloid light-chain (AL) amyloidosis, or non-Hodgkin’s lymphoma subtypes.^2^ In clinical research, the efficiency of proteasome inhibitors is typically assessed by determination of cell viability. Viability of treated cells (particularly in a clinical research context) is assessed by cell proliferation using Fluorescence-Activated Cell Sorting (FACS) assays^3,4^ in combination with detection of the formation of Endoplasmic Reticulum (ER) stress-inducing agents and pro-apoptotic proteins by Western blot (WB).^5,6^ Although these methods allow detection of specific cell-death pathways, they only deliver snapshots of data on whole cell-populations, require a large number of cells (from tens of thousands to millions) per measurement,^7,8^ and involve long and laborious multi-step sample preparation. In a clinical setting, evaluating response to multiple myeloma treatment is routinely assessed by analysis of light chains of monoclonal antibodies in serum, by immunofixation electrophoresis, microscopic or flow cytometric analysis of bone marrow aspirates and biopsies.^9,10^ These methods provide clinically relevant information on the presence of specific immunoglobulins, but they require several thousand cells, involve long laboratory procedures that prevent real-time evaluation of therapy response in patients, and only provide tumor bulk information. In contrast, a single-cell imaging technique sensitive to protein structure, requiring only a minimum number of cells and capable of rapidly assessing the heterogeneous response to proteasome inhibitors, would greatly ease treatment evaluation in individual patients using directly extracted myeloma cells. This capability is important because myeloma cells are extracted via painful biopsies, are difficult to isolate in abundance, and are challenging to maintain in culture outside the bone marrow microenvironment.^11–13^

Several imaging techniques have been employed to observe changes in protein conformation and folding states in living cells, such as single-molecule Fluorescence Resonance Energy Transfer (sm-FRET),^14^ Fluorescence Lifetime Imaging FRET (FLIM FRET),^15^ and in-cell Nuclear Magnetic Resonance (NMR).^16,17^ Although these techniques afford detection of structural and biochemical features at atomic resolution and are highly specific for tracking protein folding, they require bulky fluorescent or isotopic labels that may interfere with the study of proteins, induce cytotoxicity, and (in case of fluorophores) are prone to photobleaching.^18,19^ Additionally, Circular Dichroism has been applied to detect secondary structure conformational changes in antimicrobial peptides from random coils to α-helices during interaction with living bacterial cells;^20^ nonetheless, the analytes in the cell buffer absorb UV light leading to interference with protein signal readouts.^21^

Alternatively, vibrational spectroscopy and imaging has demonstrated label-free sensitivity to protein secondary structures, typically by detecting C=O stretching vibrations of the protein backbone in the amide I band (1700 – 1600 cm^−1^).^22,23^ For instance, Stimulated Raman Scattering (SRS) microscopy, in combination with deuterium labels or deuterated water, has been used to visualize protein aggregates in cells;^24^ however, due to low sensitivity in the amide I region,^22^ no specificity to a particular secondary structure element was demonstrated. Additionally, mid-infrared (mid-IR) spectroscopy has been proven to be a valuable tool for analysis of protein secondary structure and it is commonly used to study purified protein solutions.^25^ Nevertheless, usage of mid-IR spectroscopy/imaging for protein structure analysis in living cells is limited by the strong optical absorption of water (µ_a_: ∼742 600 M^-1^cm^-1^ at 1650 cm^−1^).^26,27^ Water opacity has been minimized in protein spectral imaging of living cells by using a cellular medium enriched by D_2_O in combination with synchrotron radiation^28^ and thin path-lengths of ≤10 µm.^29^ However, the signal-to-noise ratio (SNR) remains low due to the negative-contrast detection mechanism of conventional mid-IR spectroscopy/imaging wherein only (unabsorbed) transmitted or trans-reflected photons are detected. This represents a paradox in conventional mid-IR spectroscopy/imaging where higher optical absorption of an analyte of interest results in a lower number of photons available for its detection. Moreover, subjecting cells to high-intensity radiation (as when using synchrotron excitation) and tight spatial confinement can cause photo-damage, significantly alter the cells’ morphology, and promote deviation from their native physiological responses.^30,31^ Recently, Optical PhotoThermal InfraRed microscopy (OPTIR) was applied to image β-sheet structures and protein aggregation in primary neurons at high spatial resolution and contrast.^32–34^ However, due to the low thermo-optic coefficients of water and protein (*dn/dT*: ∼9 x 10^-5^ and ∼1.4 x 10^-4^, respectively), OPTIR applies high-intensity tightly-focused probe-beam irradiation for detection, which could induce thermal stress and photo-oxidative damage of cells over long exposure.^35,36^

Unlike conventional optical microscopy, Mid-infraRed Optoacoustic Microscopy (MiROM) is a positive-contrast modality that allows label-free spectral imaging of biomolecules in living cells by detecting optically-generated ultrasound waves that increase in intensity as optical absorption increases.^37^ We have previously demonstrated that MiROM affords higher sensitivity than Raman microscopy in the fingerprint spectral region (1800 – 900 cm^-1^) but, until now, proteins have been detected mainly by using the amide II band (1600 – 1500 cm^−1^) mainly associated with C-N stretching and N-H bending vibrations. However, the amide II spectral band does not provide sufficient specificity to protein secondary structure and, thus, cannot be used for efficient detection of intermolecular β-sheet conformational transitions.^23^

We hypothesized that the positive-contrast mechanism of MiROM in combination with partial substitution of D_2_O for H_2_O in the cell medium would yield the necessary sensitivity and specificity in the amide I region for label-free structure-specific detection of protein contrast in living-cells without the need for intense radiation and ultra-thin path lengths. Access to structure-specific protein contrast in living cells allows label-free monitoring of proteasome inhibition response in cancer therapy by exploiting the spectral features of intermolecular β-sheet structures as intrinsic biomarker of cell vitality. Thus, aiming for clinical translation, we applied MiROM to monitor intermolecular β-sheet formation upon myeloma therapy with lenalidomide (LEN) (immunomodulatory drug) and bortezomib (BTZ) (proteasome inhibitor) in an immortalized myeloma cell line (MM1.S) as well as in primary myeloma cells aspirated from patients.

Our results demonstrate that MiROM can assess patient-specific efficacy of myeloma treatment in real time at a single-cell level in small cell populations. Evaluation of myeloma therapy by direct observation of changes in protein secondary structure instead of by reduction of monoclonal proteins concentration in serum could contribute to speed up and optimize the efficacy of medical interventions. Moreover, achieving detection of treatment response at a single-cell level allows assessment of heterogeneity in primary myeloma cells, which could provide crucial therapeutic information (such as individual sensitivity to treatment) for optimizing personalized intervention.^38^

## Results

Spectroscopic analysis of the amide I region (1700 – 1600 cm^−1^) allows differentiation of protein secondary structures because the optical absorption in this spectral region is sensitive to the conformations of peptide bonds (**Fig. 1a**).^39^ First, we demonstrated MiROM’s ability to distinguish between different secondary structural motifs using proteins solutions with different proportions of α-helices and β-sheets (hemoglobin with ∼75% α-helices and concanavalin A with ∼54% β-sheets).^40^ As common practice in conventional mid-IR spectroscopy of proteins, D_2_O was used as a solvent instead of H_2_O in order to minimize optical absorption in the amide I region^41^ (see **Supplementary Fig. 1** and **Methods** section A for details). As expected, and in good agreement with measurements by Attenuated Total Reflection Fourier-Transform-InfraRed (ATR-FTIR) spectroscopy (see **Table 1**), the spectrum of hemoglobin featured a characteristic absorption band at 1650 cm^−1^ typical of α-helices (**Fig. 1b,d**),^42^ while the spectrum of concanavalin A featured a main absorption band at 1634 cm^−1^ as well as two minor side bands at 1622 and 1691 cm^−1^, all of which could be attributed to β-sheet structures (**Fig. 1c,d**).^27^ Moreover, we demonstrated MiROM’s ability to detect conformational changes in proteins by recording amide I spectra before and after heat-induced denaturation of albumin (see **Fig. 1e** and **Methods** section A). Albumin is known to undergo irreversible denaturation above 70°C and its conformational change induced by temperature has been extensively studied and characterized by mid-IR spectroscopy.^43^ In agreement with literature, we observed a peak at 1654 cm^−1^ representative of α-helices when albumin was in its native state.^43^ This peak at 1654 cm^−1^ flattens upon albumin denaturation, leading to a peak at 1612 cm^−1^. Importantly, the presence of a spectral feature at 1612 cm^-1^ is associated with the formation of intermolecular β-sheets and is a specific marker of albumin’s irreversible denaturation.^43^

**Figure 1.**
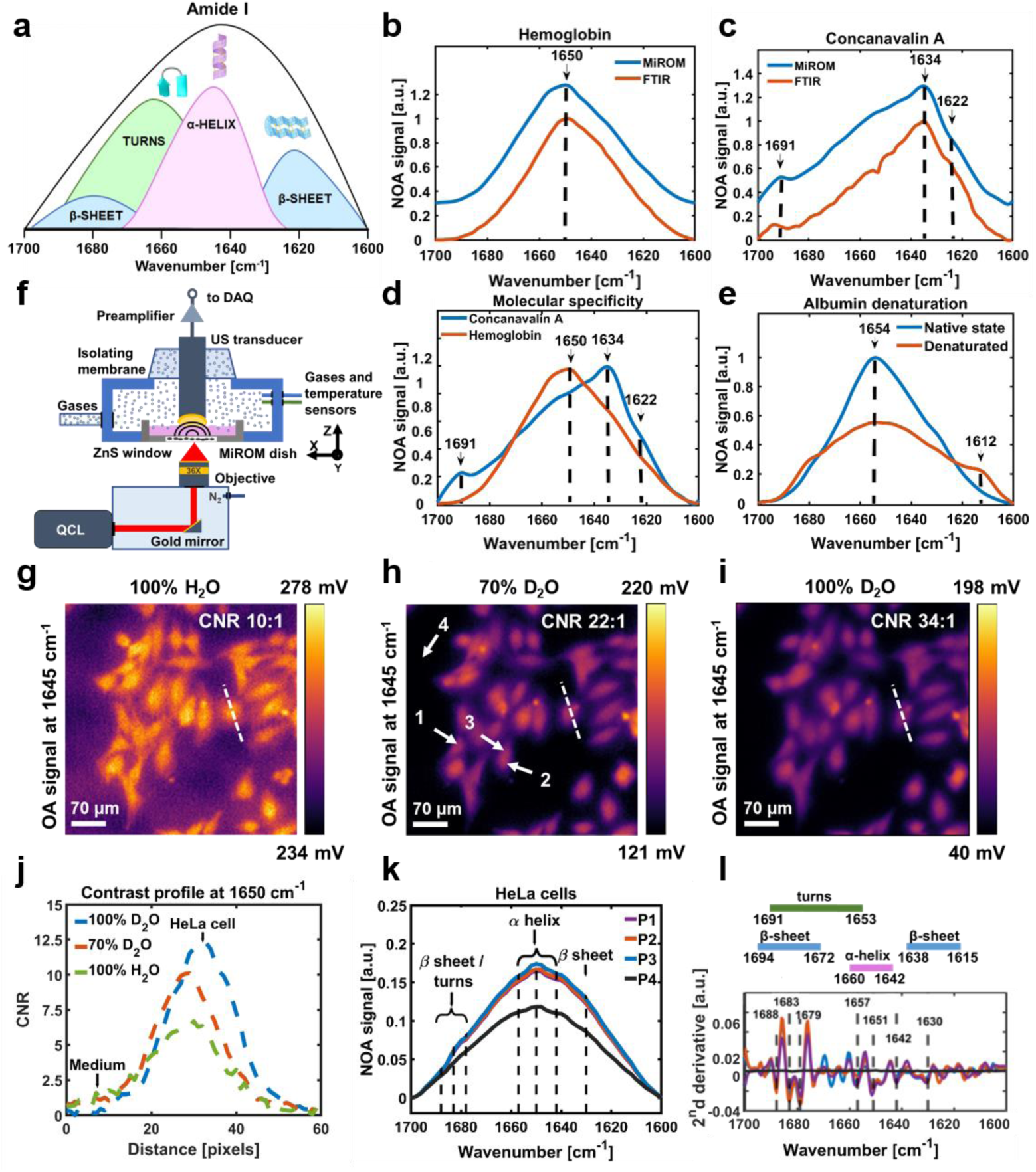
Label–free detection of the secondary structure of proteins. **a)** Illustration of the amide I Mid-Infrared (mid-IR) spectrum of proteins (1700-1600 cm^-1^) showing the absorption bands of different secondary structures. Modified from.^35^ **b)** Comparison of mid-IR absorption spectra of hemoglobin measured by Mid-iR Optoacoustic Microscopy (MiROM, blue line) and standard Attenuated Total Reflection Fourier Transform InfraRed (ATR-FTIR) spectroscopy (red line). Hemoglobin is an α-helix protein, with a characteristic band at 1650 cm^−1^.^38^ **c)** Comparison of mid-IR absorption spectra of concanavalin A measured by MiROM (blue line) and standard ATR-FTIR spectroscopy (red line). Concanavalin A is a homotetramer protein with a β-sheet structure and characteristic absorption bands at 1622, 1634 and 1691 cm^−1^.^28^ **d)** Comparison of mid-IR absorption spectra of hemoglobin (red line) and concanavalin A (blue line) measured by MiROM. **e)** Comparison of mid-IR absorption spectra of albumin in the native state (primarily α-helix, blue line) and denatured state (primarily β-sheet, red line). The vertical dashed lines indicate the α-helix absorption peak at 1654 cm^−1^ and the intermolecular β-sheet absorption peak at 1612 cm^−1^. **f)** Schematic diagram of MiROM imaging system. A tunable Quantum Cascade Laser (QCL) provides excitation for optoacoustic imaging while a focused ultrasound (US) transducer is used for signal readout. **g-i)** MiROM micrographs of live HeLa cells imaged at 1645 cm^−1^ in cell media composed of different H2O/D2O proportions. **j)** Contrast to Noise Ratio (CNR) profile of HeLa cells in (g-i). The contrast profile in 100% D2O is 3.4 times higher than the one in pure H2O, while the contrast profile in 70% D2O is 2.2 times higher than the one in pure H2O. **k)** Normalized spectra in the amide I region of live HeLa cells in 70% D2O medium acquired at the points highlighted by the arrows in (h). The vertical dashed lines indicate the turn, α-helix, and β-sheet secondary structure absorption peaks (see **Table 2**). **l)** Second derivative plot of spectra from (k). The scheme above the plot details the values corresponding to the different secondary structures according to the literature.^43^ Three independent experiments (n=3). OA – Optoacoustic. NOA – Normalized Optoacoustic.

**Table 1.**
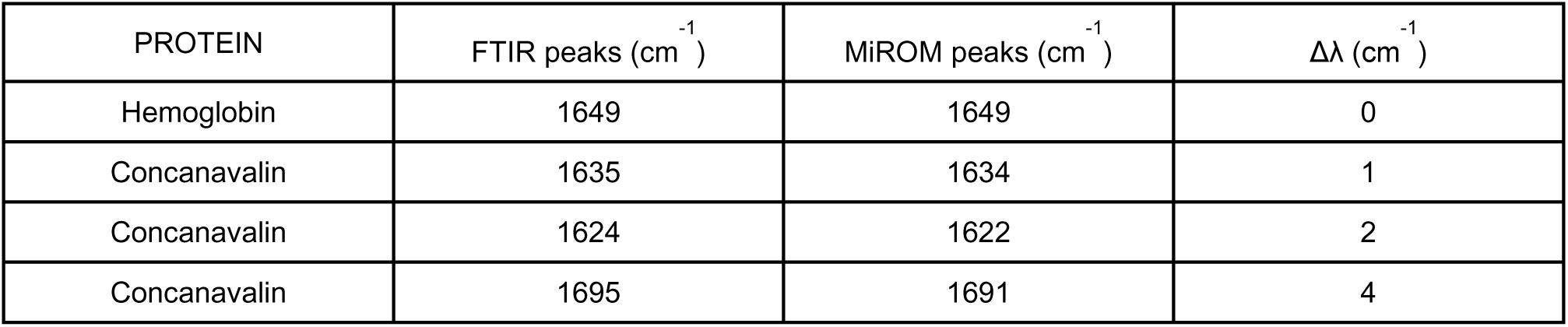
Relevant absorption peaks of hemoglobin, concanvalin by Attenuated Total Reflection Fourier-Transform-InfraRed (ATR-FTIR) spectroscopy and by Mid-infraRed Optoacoustic Microscopy (MiROM). The difference (Δλ) between ATR-FTIR and MiROM’s values is indicated in the third column.

| PROTEIN | FTIR peaks ($\text{cm}^{-1}$) | MiROM peaks ($\text{cm}^{-1}$) | $\Delta\lambda$ ($\text{cm}^{-1}$) |
| --- | --- | --- | --- |
| Hemoglobin | 1649 | 1649 | 0 |
| Concanavalin | 1635 | 1634 | 1 |
| Concanavalin | 1624 | 1622 | 2 |
| Concanavalin | 1695 | 1691 | 4 |

**Table 2.**
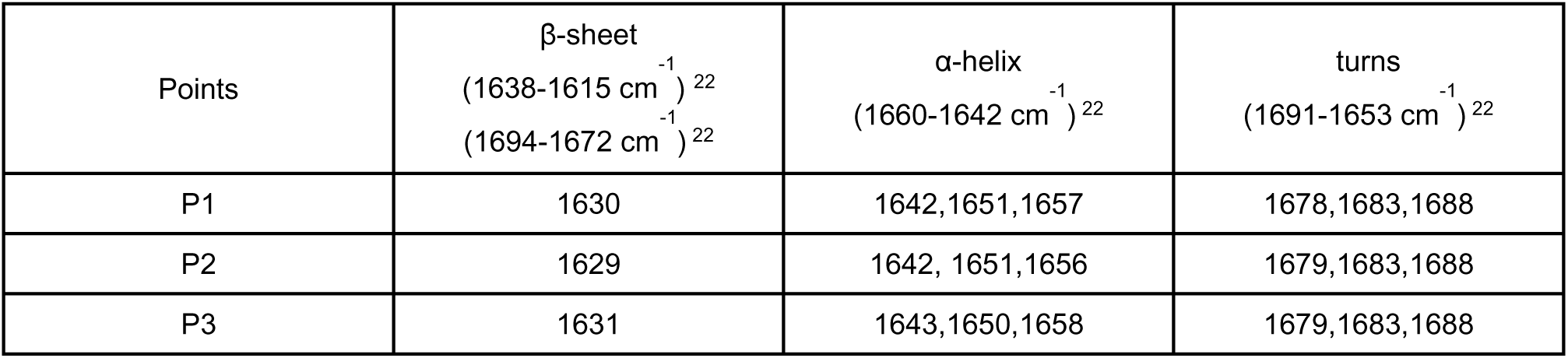
Comparison between the peak values of different secondary structures detected in live HeLa cells by by Mid-infraRed Optoacoustic Microscopy (MiROM) and the values reported in literature.^22^

| Points | β-sheet<br>(1638-1615 $\text{cm}^{-1}$)<sup>22</sup><br>(1694-1672 $\text{cm}^{-1}$)<sup>22</sup> | α-helix<br>(1660-1642 $\text{cm}^{-1}$)<sup>22</sup> | turns<br>(1691-1653 $\text{cm}^{-1}$)<sup>22</sup> |
| --- | --- | --- | --- |
| P1 | 1630 | 1642,1651,1657 | 1678,1683,1688 |
| P2 | 1629 | 1642, 1651,1656 | 1679,1683,1688 |
| P3 | 1631 | 1643,1650,1658 | 1679,1683,1688 |

We next applied MiROM to analyze the amide I region of HeLa cells in D_2_O/H_2_O medium in order to investigate its sensitivity to protein secondary structure in living cells. To guarantee a controlled environment for the cells’ growth, we outfitted MiROM with a mini-incubator, which maintains optimal humidity, and flux of oxygen and CO_2_ (**Fig. 1f**). HeLa cells were first imaged at 1645 cm^-1^ in cell media composed of different D_2_O/H_2_O percentages; 0/100% (**Fig. 1g**), 70/30% (**Fig. 1h**), and 100/0% (**Fig. 1i**), respectively. Notably, we observed that MiROM could image HeLa cells even in 100% H_2_O, albeit with a lower contrast than in media with higher D_2_O percentages (**Fig. 1g-j** and **Supplementary Fig. 2**). Using 100% D_2_O afforded an enhancement in contrast-to-noise ratio (CNR, see **Methods** for details) up to 3.4 times compared to using pure H_2_O, while using 70% D_2_O afforded an enhancement of 2.2 times (CNR 34:1 in 100% of D_2_O, **Fig. 1i**; CNR 22:1 in 70% D_2_O, **Fig. 1h**; CNR 10:1 in pure H_2_O, **Fig. 1g**; see **Supplementary Fig. 2** and **Methods** section B). As shown in **Fig. 1g-i**, higher contrast was obtained when imaging HeLa cells in 100% D_2_O as compared to cell media with lower D_2_O percentages. We demonstrated cell viabilities up to 93.8% in HeLa cells exposed to 70% D_2_O and up to 85.2% when HeLa cells were exposed to 90% D_2_O over 24 hours exposure time (see **Supplementary Fig. 3** and **Methods** section B for details). However, long exposure to D_2_O can cause cell toxicity, therefore we reduced exposure time to 2 hours maximum in all experiments. MiROM was then used to extract spectra from several intracellular locations in live HeLa cells (white arrows in **Fig. 1h**). Inspection of the spectral content at these locations revealed several features associated with protein secondary structure in the amide I region, such as α-helices (1660 – 1642 cm^−1^),^23^ β-sheets (1638 – 1615 cm^−1^ and 1694 – 1672 cm^−1^),^23^ and turns (1691 – 1653 cm^−1^)^23^ (see **Fig. 1k, l** and **Table 2**). As expected and in good agreement with observations of lyophilized cells by FTIR spectroscopy combined with synchrotron excitation,^44^ the spectral content obtained with MiROM at different cell locations indicates a homogeneous distribution of protein structures (Δλ=2 cm^-1^ maximum spectral deviation, **Fig. 1l**).

We then set out to apply MiROM to detect conformational transitions of proteins in a cancer cell model (namely: multiple myeloma), where accumulation of misfolded proteins marks a central therapeutic mechanism for approved drugs. The myeloma cell line, MM1.S, produces high amounts of dysfunctional immunoglobulins (paraproteins); treatment with the immunomodulatory drug LEN (IMiD) and the 26S proteasome inhibitor BTZ results in accumulation and aggregation of intracellular misfolded proteins, ultimately leading to cell apoptosis (**Fig. 2a**).^5,45–48^ Misfolded proteins are rich in intermolecular β-sheet structures,^4^ which MiROM can detect using the proteins’ characteristic spectral features in the amide I band. Therefore, we propose detection of intermolecular β-sheet structures as a biomarker for treatment response. For demonstration, individual MM1.S cells in 100% D_2_O medium were arbitrarily selected from MiROM micrographs—at 1650 cm^-1^ and 500 x 500 µm^2^ Field-of-View (FOV)—for spectral analysis in the range between 1700 and 1600 cm^-1^ (**Fig. 2b**). The spectral analysis was conducted, prior to treatment and then at different time points after LEN/BTZ treatment, drugs commonly used in multiple myeloma (MM) therapy (see **Methods** section C for details).^49,50^ Myeloma cells were maintained in 100% D_2_O medium only during MiROM measurements, for a maximum of 2 hours. Spectra, acquired from cells by MiROM after 96 hours of LEN/BTZ treatment, exhibited broadening of the amide I band towards frequencies characteristic of β-sheets (red line, **Supplementary Fig. 4a**). Differential spectra (spectral difference before and after treatment, see **Methods**) further revealed the formation of a prominent spectral feature at ∼1620 cm^-1^ (**Fig. 2c** shows representative example from five independent experiments). This spectral feature is attributed to antiparallel intermolecular β-sheets^39,51^ and suggested the presence of misfolded protein aggregates. A similar spectral band, indicating protein aggregation, was found (in a separate study by other authors) in amyloid fibrils containing β-sheets in phantoms and in human tissues.^23,39,52^ As a control, cells neither treated with LEN nor with BTZ were spectrally analyzed with MiROM in the same way (**Fig. 2d** and **Supplementary Fig. 4**b). No spectral changes were observed in untreated cells over time (**Fig. 2c** and **Supplementary Fig. 4b**). Spectral analysis at treatment points earlier than 96 hours reveals the appearance of the intermolecular β-sheet band after 84 hours of treatment (**Supplementary Fig. 5**), with the band becoming more intense and pronounced 8 hours after (at 92, **Fig. 2e**). No intermolecular β-sheet band was observed at earlier treatment times (at 78 hours, **Supplementary Fig. 5**). Interestingly when studying the individual effects of LEN^5^ or BTZ^5,53^ treatment, we observed a similar formation of β-sheet intermolecular secondary structures in myeloma cells as a treatment response. This is shown in **Fig. 2f,g**, where a prominent band at 1620 cm^-1^ appeared after MM1.S cells were subjected to 25 μM LEN (treatment response expressed in 50% of cells after 96 hours and in 100% of cells after 144 hours) or to 100 nM BTZ (treatment response expressed in 80% of cells after 18 hours, see **Supplementary Fig. 6** and 7, and **Method** section C). MiROM’s detection of intermolecular β-sheet formation at 1620 cm^-1^ upon LEN treatment alone suggests that (similarly to BTZ) LEN might also result in misfolded protein accumulation—advocating for combinatory LEN/BTZ treatment as a strategy to increase efficacy.^5^

**Figure 2.**
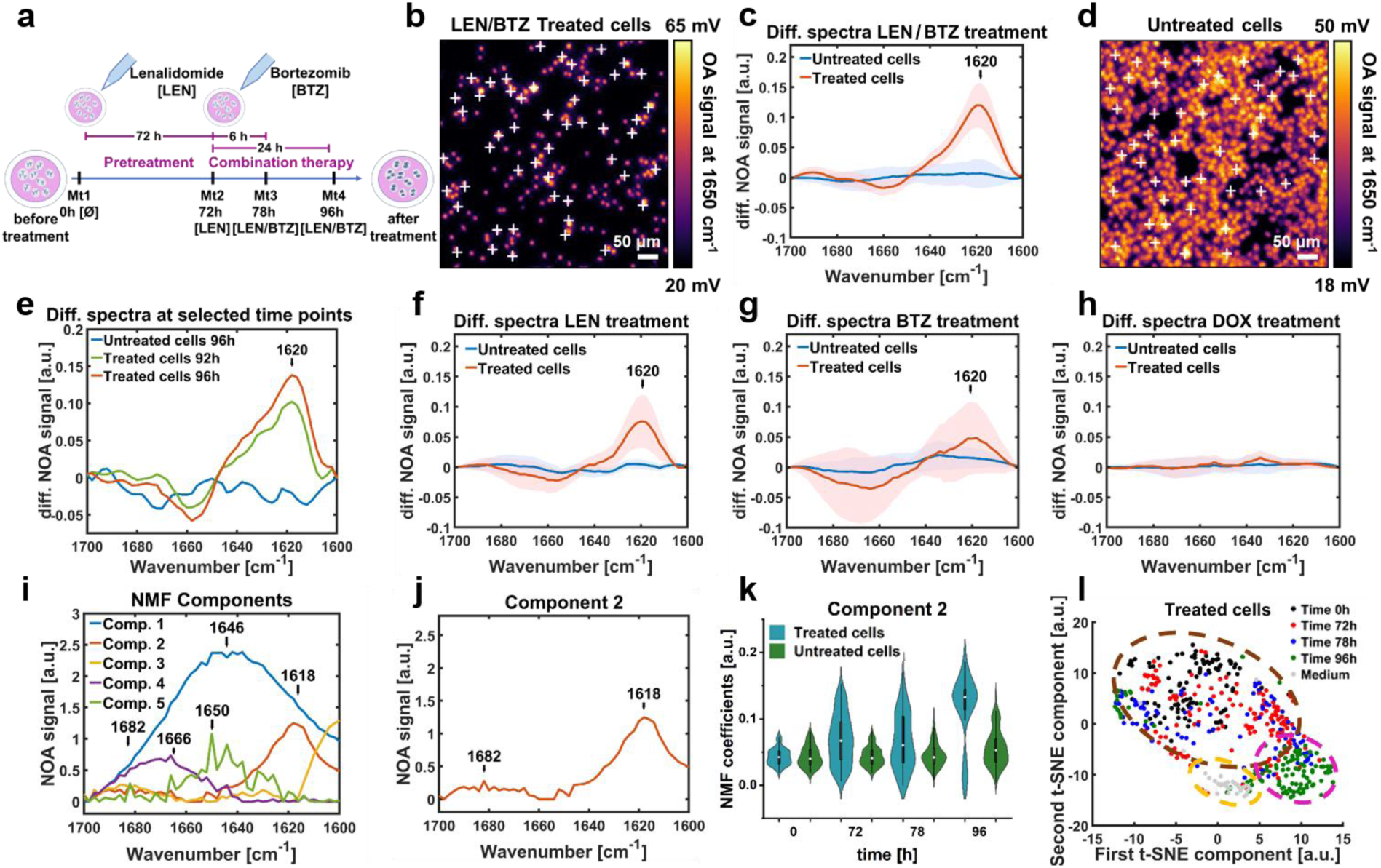
Monitoring protein misfolding in myeloma cells during synergic and individual lenalidomide (LEN) and bortezomib (BTZ) treatment. **a)** Schematic diagram of LEN and BTZ treatment in myeloma cells. **b)** Multiple myeloma cancer cells (MM1.S) pretreated with 10 μM of LEN for 72 hours and, subsequently, treated with 100 nM of BTZ for 24 hours imaged at 1650 cm^-1^. The image was acquired after 96 hours of treatment (Mt4). **c)** Differential spectra of the LEN/BTZ treated myeloma cells shown in b (red line), and the untreated myeloma cells shown in d (blue line) at 96 hours. The spectrum in red shows a band at 1620 cm^-1^, assigned to an intermolecular β-sheet of misfolded proteins which was not found in untreated cells. **d)** Untreated myeloma cells imaged at 1650 cm^-1^ after being in culture for 96 hours. **e)** Differential spectra of LEN/BTZ treated MM1.S cells at different time points (green: 92 hours, red: 96 hours). The differential spectrum of untreated cells is shown in blue. MiROM can detect the effect of proteasome inhibition after 92 hours of LEN and BTZ combined treatment (n=5 independent experiments). **F)** Differential spectra of MM1.S cells treated with 25 μM of LEN for 144 hours compared to untreated cells maintained in culture for 144 hours. LEN-treated cells show formation of the intermolecular β-sheet structure, which is not present in the untreated cells (n=3 independent experiments). **G)** Differential spectra of MM1.S cells treated with 100 nM of BTZ for 18 hours compared to untreated cells maintained in culture for 18 hours. BTZ-treated cells show formation of the intermolecular β-sheet structure, which is not present in the untreated cells (n=3 independent experiments). **h)** Differential spectra of MM1.S cells treated with 1 μM of doxorubicin (DOX) for 2 hours compared to untreated cells maintained in culture for 2 hours. The intermolecular β-sheet structure is not visible in either differential spectra (untreated and DOX-treated cells, n=3 independent experiments). **i)** Non-negative Matrix Factorization (NMF) components extracted from spectral data (n=700 spectra). **j)** Component 2 represents intermolecular β-sheet secondary structures (1682-1618 cm^-1^). **k)** Violin plots from kernel density estimate the time evolution coefficients of NMF component 2 in LEN/BTZ treated cells and in untreated cells. Component 2 increased by 174% during the treatment. **l)** A t-Distributed Stochastic Neighbor Embedding (t-SNE) map representing the distribution of the 5 components identified in LEN/BTZ treated myeloma cells. Spectra acquired from the cell medium and spectra acquired from treated cells after 96 hours of treatment are clustered together. OA – Optoacoustic. NOA – Normalized Optoacoustic.

In order to assess whether the observed changes in the protein secondary structure were specific for BTZ or LEN treatment, MM1.S cells were treated with doxorubicin (DOX)—a chemotherapeutic drug that causes cell death by damaging single and double strand DNA without inducing accumulation and aggregation of misfolded proteins (**Fig. 2h**, **Supplementary Fig. 8** and **Method** section C).^54^ Indeed, the comparison between the differential spectra of cells with or without DOX treatment revealed the absence of the β-sheet intermolecular band, providing evidence of MiROM’s specificity to the mechanism of action for LEN and BTZ (**Fig. 2h**).

In order to obtain a more detailed picture of spectral changes during MM1.S treatment (than with the differential spectra analysis; **Fig. 2c**), the acquired (LEN/BTZ treated and untreated) single-cell spectra (∼700 spectra from five independent experiments) were analyzed by regularized Non-negative Matrix Factorization (NMF). The number of spectral components necessary to suitably represent the data set was estimated via Principal Component Analysis (PCA) (see **Methods** section C and **Supplementary Fig. 9a**). **Figure 2i** shows the five main components obtained by NMF analysis of spectra from untreated and LEN/BTZ treated cells. Component 1, centered at 1646 cm^-1^, describes a common feature within the spectra acquired associated to the cell medium. Component 2, at 1618 cm^-1^ is associated with the presence of intermolecular β-sheets (see **Fig. 1a**). Component 3 describes the emission profile of the laser (**Supplementary Fig. 9b**). Component 4, at 1666 cm^−1^, and component 5, at 1650 cm^-1^, can be associated with the presence of α-helix structures (see **Fig. 1a**) and are influenced by the exchange between the hydrogen in the atmosphere and the deuterium in the medium (see **Supplementary Fig. 9c**). In **Fig. 2j**, component 2 shows a second band at 1682 cm^-1^ (characteristic of intermolecular β-sheet structures^39^), which was not clearly visible from the differential spectra analysis. According to literature, the band at 1682 cm^-1^ is weaker than the one at 1620 cm^-1,39^ and therefore more difficult to detect by MiROM alone without the application of NMF analysis. In **Fig. 2k**, violin plots show the time evolution of component 2 for LEN/BTZ treated and untreated cells. In treated cells, component 2 increases by 174% after 96 hours of treatment with LEN/BTZ (violin plots in blue, **Fig. 2k**), confirming the increase of intermolecular β-sheet secondary structures as observed by the differential spectral analysis (**Fig. 2c**). The time evolution of component 2 describes the trend of both intermolecular β-sheet bands (1689 and 1620 cm^-1^). We noticed (**Supplementary Fig. 9**f, g) that the decreasing dynamics of component 4 (∼33%) and 5 (∼25%), suppresses the intensity increase at the 1689 cm^-1^ band, which might explain why 1689 cm^-1^ is not visible in the differential spectrum (**Fig. 2c**). Moreover, the decreasing dynamics of components 4 and 5 generates a dip at ∼1650 cm^-1^ in the differential spectra, suggesting a reduction in α-helix structures. As expected, in untreated cells, the time evolution of component 2 does not show substantial changes (violin plots in green, **Fig. 2k**), confirming the absence of intermolecular β-sheet formation. This result is in agreement with our differential spectral analysis (**Fig. 2c**) that shows no change in the spectra of untreated cells. A stable trend for component 5 in untreated cells suggests retaining of α-helix structures in the cells’ proteins; for components 1-3, the time evolution changes are negligible (see **Supplementary Fig. 9**d-g for more details). For better visualization, PCA and NMF analysis of the measured spectra yielded a two dimensional t-distributed Stochastic Neighbor Embedding (2D t-SNE) map illustrating the contributions of five molecular spectral components to each spectrum (see **Methods** section C). The t-SNE map of LEN/BTZ treated cells in **Fig. 2l** shows three separate groupings: (1) spectra acquired from 0 to 78 hours (brown circle), (2) spectra acquired after 96 hours (magenta circle), and (3) spectra of the medium (orange circle). The grouping of points on the t-SNE maps reflects similarities in protein secondary structure. This means that the spectra taken after 96 hours of combinatory treatment with LEN/BTZ were spectrally different to those obtained at any earlier points. In a similar fashion, the t-SNE map of untreated cells (**Supplementary Fig. 9h**) shows only 2 groups: one composed of cells spectra acquired at all-time points (magenta circle) and a second group composed of spectra of the media (blue circle); these groupings indicate that there are no changes in secondary structure and are in agreement with the differential spectral analysis, **Fig. 2c**.

The presence of the spectral band at 1620 cm^-1^, obtained by differential spectra (**Fig. 2c**) as well as by NMF analysis (**Fig. 2i-k**), clearly indicates formation of intermolecular β-sheets in proteins of MM1.S cells when treated with LEN or BTZ individually or in combination; thus, demonstrating MiROM’s ability to identify intrinsic hallmarks of misfolded proteins and aggresome formation, and treatment response without the use of labels or a large number of cells. Our observations with MiROM are in agreement with observations (by other authors) using immunoblots, apoptosis and MTT assays, where apoptotic cell death of MM1.S cells occurred in response to LEN or BTZ treatment.^5,45–49,55^

After demonstrating sensitivity to misfolded protein formation in immortalized cell lines during LEN or BTZ treatment individually or in combination as a first step towards clinical translation, we applied MiROM to analyze protein misfolding in CD138^+^ purified primary myeloma cells from newly diagnosed myeloma patients (10 patients measured independently). Following the same procedure applied to MM1.S cells, the patients’ cells were measured in 100% D_2_O medium for circa 2 hours. **Figure 3a-f** exemplifies results from one representative LEN/BTZ-sensitive myeloma patient (with data from the 10 patients summarized in **Table 3**). Individual cells were arbitrarily selected from MiROM micrographs (at 1650 cm^-1^, 500 x 500 μm^2^ FOV, **Fig. 3a**) for spectral analysis (in the amide I region) at different time points: namely, before adding any drug (time 0h), 48 hours after LEN administration (before adding BTZ), and 72 hours after LEN and BTZ combinatory treatment (see **Method** section D). As a control, untreated myeloma cells biopsied from the same patient were measured in the same way (**Fig. 3b**). Consistent with our results in treated MM1.S cells (**Fig 2c** and **Supplementary Fig. 4a**), spectral analysis of treated patient cells revealed spectral broadening towards lower frequencies (**Supplementary Fig. 10a**) and the presence of the band at 1620 cm^-1^, which is characteristic of intermolecular β-sheet structures of misfolded proteins (**Fig. 3c**). As expected, the spectra of untreated cells did not show a prominent band of intermolecular β-sheet structure (**Fig. 3c** and **Supplementary Fig. 10b**), supporting specificity of 1620 cm^-1^ as an intrinsic spectral marker of treatment response. Moreover, PCA and NMF analysis of patient cells’ spectra identified 5 main components similar to the ones obtained in the analysis of the MM1.S cell line (**Fig. 2i**); although in a different order of appearance (see **Fig. 3d** and **Supplementary Fig. 11** for details). In particular, in LEN/BTZ-treated patients’ cells, component 4 at 1620 cm^-1^ (indicative of intermolecular β-sheet structures) showed an intensity time evolution that increased (after 72 hours of combined treatment) by 44% (**Fig. 3e**), component 5 at 1656 cm^-1^ (α-helix indicator) showed decreased intensity by 52% (**Supplementary Fig. 11e**), while the change over time in the remaining components (1-3) was found to be negligible (**Supplementary Fig. 11**c-e). Importantly, untreated cells showed negligible change over time for all the 5 components, suggesting unchanged protein secondary structures and supporting our previous observations from differential spectra analysis of untreated MM1.S cells (**Supplementary Fig. 11**c**-f**). Additionally, the 2D t-SNE maps of the five molecular spectral components’ coefficients resulted in separated clusters at the different time points after LEN/BTZ treatment (**Fig. 3f**). Furthermore, there was a unique cluster for the spectral components of untreated cells (**Supplementary Fig. 11b**), indicating that the spectral components of treated cells changed during treatment while remaining stable in untreated cells.

**Figure 3.**
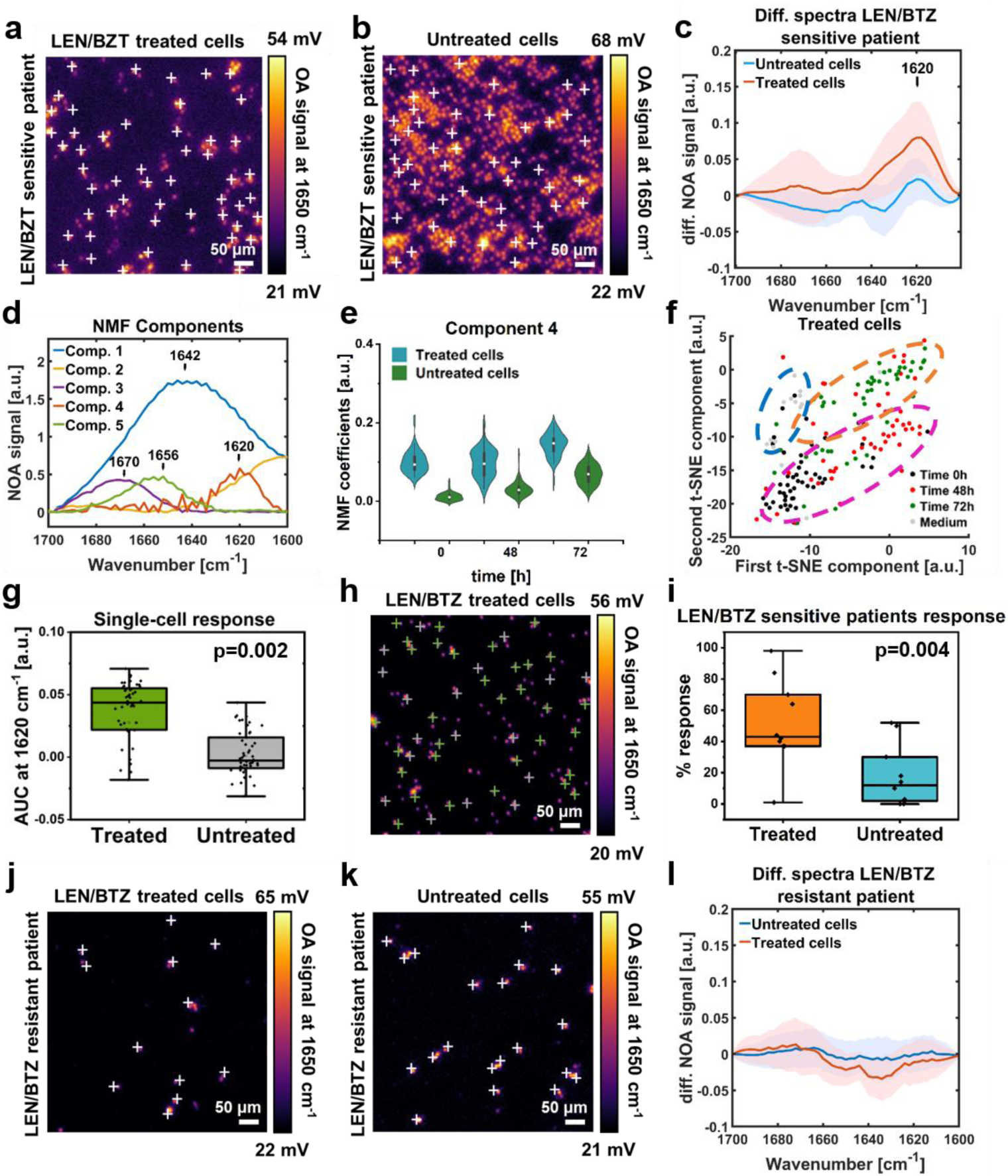
Monitoring protein misfolding of myeloma cells sensitive or resistant to lenalidomide (LEN) and bortezomib (BTZ) in patients. **a,b)** Myeloma cells extracted from the bone marrow of a LEN and BTZ sensitive patient were plated in two separate dishes (see **Methods** for details). One dish (a) was imaged at 1650 cm^-1^ after pretreatment for 48 hours with 10 μM of LEN and then treatment for 24 hours with 100nm of BTZ. The second dish (b) was used as a control (untreated) and imaged at 1650 cm^-1^ after being in culture for 72 hours. **c)** Comparison between the differential spectra of LEN/BTZ treated (in red) and untreated (in blue) myeloma cells after 72 hours. LEN/BTZ treated cells show the absorption band of an intermolecular β-sheet structure at 1620 cm^-1^ (n=10 patients). **d)** Non-negative Matrix Factorization (NMF) components extracted from spectral data in (c) (n=300 spectra). **e)** Violin plots from kernel density estimate the time evolution coefficients of NMF component 4 in LEN/BTZ treated cells and untreated cells. Component 4 increased by 44% during the treatment. **f)** A t-Distributed Stochastic Neighbor Embedding (t-SNE) map representing the distribution of the 5 components identified in LEN/BTZ treated myeloma cells. **g)** Boxplot representing the area under the curve (AUC) for the band at 1638 – 1615 cm^-1^of LEN/BTZ treated and untreated cells extracted from one LEN and BTZ sensitive patient. **h)** 32 single cells identified by the AUC analysis show the intermolecular β-sheet band (green cross); while 18 cells do not show the β-sheet band (grey cross). **i)** Boxplots representing the percentage response (%) of LEN/BTZ treated and untreated cells analyzed from 10 independent patients sensitive to LEN and BTZ. **j, k)** LEN/BTZ treated (j) and untreated (k) myeloma cells extracted from the bone marrow of a LEN and BTZ resistant patient. P values from paired sample *t*-test. **l)** Comparison between differential spectra of LEN/BTZ treated and untreated myeloma cells in (j,k). Both, LEN/BTZ treated and untreated cells show no spectral band in the region of the intermolecular β-sheet structure (1638 – 1615 cm^-1^), denoting the absence of misfolded protein formation under proteasome inhibition (n=2 patients). OA – Optoacoustic. NOA – Normalized Optoacoustic.

**Table 3.** Percentage response (%) of untreated and lenalidomide (LEN) and bortezomib (BTZ) treated cells, shown per patient and calculated counting the number of single cells that show formation of β-sheet intermolecular structure during spectral imaging with Mid-infraRed Optoacoustic Microscopy (MiROM).

| Patient | LEN/BTZ treated cells |  |  | Untreated cells |  |  |
| --- | --- | --- | --- | --- | --- | --- |
| First diagnosis | β-sheet band | No β-sheet band | % response | β-sheet band | No β-sheet band | % response |
| 1 | 47 cells | 9 cells | 84% | 27 cells | 25 cells | 52% |
| 2 | 20 cells | 30 cells | 40% | 15 cells | 35 cells | 30% |
| 3 | 22 cells | 28 cells | 44% | 0 cells | 50 cells | 0% |
| 4 | 32 cells | 18 cells | 64% | 1 cell | 49 cells | 2% |
| 5 | 49 cells | 1 cells | 98% | 5 cells | 45 cells | 10% |
| 6 | 18 cells | 31 cells | 37% | 0 cells | 50 cells | 0% |
| 7 | 22 cells | 31 cells | 42% | 7 cells | 43 cells | 14% |
| 8 | 2 cells | 48 cells | 1% | 9 cells | 41 cells | 18% |
| 9 | 15 cells | 26 cells | 37% | 1 cell | 37 cells | 3% |
| 10 | 37 cells | 16 cells | 70% | 28 cells | 28 cells | 50% |
| **Total** |  |  | **55%** |  |  | **24%** |
| Refractory | β-sheet band | No β-sheet band | % response | β-sheet band | No β-sheet band | % response |
| 1 | 0 cells | 54 cells | 0% | 1 cell | 51 cells | 2% |
| 2 | 4 cells | 46 cells | 8% | 4 cells | 46 cells | 8% |
| **Total** |  |  | **1%** |  |  | **1%** |

In order to take advantage of MiROM’s specificity to intermolecular β-sheet formation as well as its ability for single-cell analysis, we studied the heterogeneity of therapy efficacy in biopsied cells for assessment of treatment response in individual patients. For this purpose, we defined a metric to determine single-cell response based on the spectral content (i.e., Area Under the Curve, AUC) in the range specific for intermolecular β-sheet structures (1638 – 1615 cm^-1^) (**Methods** section D and **Supplementary** Fig.12 **a-e** for details). Cells with AUC bigger than the threshold value (defined by AUC of untreated cells at 0h) have spectral features indicating development of intermolecular β-sheets and these cells were classified as responsive to treatment. While cells with AUCs smaller or equal than the threshold value did not show the relevant band and were, thus, classified as unresponsive. The percentage of responding cells, including details on the number of unresponsive or responsive cells, are summarized in **Table 3**. As exemplified by the boxplots in **Fig. 3g** (from one representative patient), LEN/BTZ-treated cells have higher AUC values compared to untreated cells (boxplots for all patients shown in **Supplementary** Fig.13). **Figure 3h** shows the distribution of the cells defined as responsive or unresponsive for one representative patient; 32 responsive cells (white marks) and 18 unresponsive cells (green marks) yielding a positive treatment response in 64% of the total cells. The mean positive treatment response calculated in all the patients analyzed was 52% overall. The mean response in untreated cells showed a notably lower value at 18%.

Finally, we applied MiROM to assess therapy response in relapsed/refractory (r/r) patients previously exposed to both LEN and BTZ treatment (**Figure 3j-l**). Patients with relapsed/refractory myeloma refers to individuals who either did not respond to the treatment at all or experienced a more aggressive recurrence of cancer after a period of remission. In those r/r MM patients, no, or negligible, formation of intermolecular β-sheet structures was expected. Comparable to observations in control experiments (i.e., in untreated cells as well as in cells treated with DOX), the differential spectra of cells from refractory patients treated with LEN/BTZ (**Fig. 3l**) reveals absence of the band indicating intermolecular β-sheet structures at 1620 cm^-1^. Thus, MiROM demonstrated its ability to evaluate not only the positive response of LEN/BTZ-sensitive patients, but also the negative response of LEN/BTZ-resistant patients. In accordance, boxplots of AUCs, from spectra of LEN/BTZ treated or untreated cells from r/r patients, do not show significant differences (i.e., the responses in both LEN/BTZ treated and untreated cells were below 1%, see **Supplementary Fig. 14** and **Table 3**). Consequently, our results on unresponsive patients provides additional support that MiROM is able to specifically assess LEN and BTZ therapy efficacy in clinical samples via detection of intermolecular β-sheet structures, and its potential in terms of response assessment.

## Discussion

Conventional optical microscopy techniques (in particular, those operating in the visible spectral region) lack the sensitivity and specificity for label-free detection of protein secondary structures in living cells. Here, we demonstrated that MiROM achieves label-free protein-structure specific sensitivity in living cells. Instead of applying exogenous markers, MiROM uses the intrinsic spectroscopic signatures of protein secondary structure in the amide I band (1700 — 1600 cm^-1^) as endogenous source of contrast. This unprecedented ability enabled for the first time label-free detection of intermolecular β-sheet formation in primary myeloma cells and MM1.S cells treated with LEN or BTZ individually or in combination. Intermolecular β-sheet formation is a hallmark of misfolded protein accumulation and aggresome formation, which we used as an indicator of cell apoptosis.

Unlike conventional clinical techniques for evaluating therapy response, i.e., serum light chain and protein electrophoresis, immunofixation or flow cytometry of bone marrow biopsies, MiROM evaluations can be performed in real time directly after purification of myeloma cells from the bone marrow. The assessment of myeloma therapy response by observation of intermolecular β-sheet formation instead of detection of reduced monoclonal protein quantities in serum or bone marrow aspirates (with a latency period of 3 to 4 weeks) could accelerate prediction of myeloma therapy efficacy and outcome. Moreover, MiROM assessment is unlike WB and FACS assays currently used in clinical research to study cell viability^7,8^ because MiROM requires only a minimum amount of cells and not tens of thousands to millions of cells per measurement point (for example, FACS requires between 30,000 and 50,000 cells and WB requires 1 million cells).^7,8,56^ Additionally, MiROM can longitudinally detect drug-induced apoptosis in real time unlike FACS and WB, which only provides a snapshot of bulk information. Typically, in MM patients, the amount of myeloma cells obtained after purification of a 10 mL liquid biopsy varies between 10,000 to 2 million cells. Therefore, the number of cells collected from each biopsy is frequently insufficient for determination of cell viability after therapy when using WB and FACS assays; this is especially challenging in longitudinal studies where several measurement points are needed to assess drug performance over time. Although additional bone marrow aspirations can be performed to increase cell count from one patient, bone marrow aspiration is associated with stinging and sharp pain, causing discomfort to patients. To increase the cell yield, novel purification methods (such as immunomagnetic bead depletion) have been applied to obtain up to 95% of the myeloma cells.^12^ Nonetheless, the number of purified cells could still be insufficient for performing longitudinal studies of cell viability. Additionally, primary myeloma cell culture and expansion is challenging or even impossible; therefore, this strategy to generate sufficient cultured cells for FACS or WB is not a feasible alternative.^13^ Thus, going beyond conventional methods, MiROM offers a powerful alternative to evaluate therapy efficacy in patients’ myeloma cells—allowing us to monitor for the first time treatment response to LEN/BTZ in primary patient samples without the use of external labels. The ability to detect apoptotic effects in a small number of cells opens up the possibility of longitudinal assessment of multiple drugs in parallel using cells from the same biopsy allowing prediction of therapy response for personalized myeloma treatment. In clinical research, evaluation of treatment in a small amount of patient cells using MiROM—as opposed to conventional methods that require a large number of primary myeloma cells—^13^ may reduce tedious and painful procedures for the patient. Indeed, MiROM demonstrated to be capable to discern between LEN/BTZ sensitive and LEN/BTZ resistant patients by detecting the formation or absence of misfolded protein structures in the amide I region after myeloma treatment. Moreover, achieving detection of therapy response to a single-cell level could allow MiROM to improve understanding of intratumoral heterogeneity by identification of cell subpopulations responsible for causing treatment failure due to drug resistance.^38^

MiROM is a positive-contrast imaging modality that attains sensitivities in the amide I region superior to conventional mid-IR microscopy and spectroscopy methods, which are predominantly negative contrast.^37^ This unique characteristic allows MiROM to overcome the limitations encountered by conventional mid-IR spectroscopy and imaging, which can only achieve the detection of protein secondary structures in living cells with the use of high power synchrotron radiation and ultra-narrow path-lengths that perturb cells’ physiology.^29,57^ The current sensitivity of MiROM allows detection of signal changes as low as 1% in the amide I region. In this study, we observed a maximum signal change up to 15% in single cells after 72 hours of LEN and 24 hours of BTZ treatment (i.e., after 96 hours of combined action) with an average change of around 8%. Further enhancing MiROM’s sensitivity—for example, by using narrower laser pulse durations to increase optoacoustic efficiency (e.g., 5 ns instead of 20 ns as used here)—would allow detecting signal changes smaller than 1%, thus resulting in earlier detection of therapeutic effects. Additionally, increasing imaging and spectral acquisition speed will allow analysis in larger cell populations (beyond the 500 x 500 µm^2^ FOV studied here), thus taking full advantage of patient cell extractions and increasing the accuracy of treatment assessment.

Finally, the herein demonstrated ability to detect intrinsic protein misfolding and aggregation during MM treatment could also become key in studying the therapy efficiency in other diseases treated with proteasome inhibitors (for example, mantle cell lymphoma and AL amyloidosis). Additionally, MiROM could also be relevant for studying protein dynamics in other protein misfolding diseases (such as Alzheimer’s or Parkinson’s disease) as well as in enabling efficient drug screening in related preclinical studies. Thus, requiring minimal sample preparation and being cost-effective, MiROM could contribute to personalized medicine by supporting identification of the most efficient therapy, sparing time and minimizing errors in optimizing patient-specific treatment.

## Methods

### Methods section A – protein solutions

To investigate the ability of MiROM to distinguish between proteins with different secondary structures, two representative proteins (hemoglobin and concanavalin A) were analysed. The samples were prepared and measured as follows: a 10 mg/mL D_2_O solution of hemoglobin was prepared by dissolving 500 mg of hemoglobin powder (HiMedia Laboratories) in 50 mL of D_2_O; a 10 mg/mL D_2_O solution of concanavalin A was prepared dissolving 200 mg of concanavalin A (Sigma-Aldrich) in 20 mL of D_2_O phosphate buffer. To investigate the ability of MiROM to monitor conformational changes in the secondary structure of proteins, we observed the effect of heat-denaturation on a D_2_O solution of albumin. A solution of 50g/L D_2_O albumin was prepared by dissolving 125 mg of albumin (Carl Roth) in 25 mL of D_2_O water. The solution was imaged spectrally at different temperatures starting from room temperature (25 °C), where albumin has a native α-helix structure, until above 80°C, where its structure is completely denatured into an intermolecular β-sheet conformation. The heating denaturation process was performed using a stage heating system and the temperature was monitored with a thermometer. All protein solutions were filtered and measured on a custom-made mid-IR dish with a ZnSe window using carbon tape (SPI Suppliers, PA, USA) as a spectral reference for correcting the systematic variation due to factors such as the mid-IR output spectrum, optical components absorption spectrum…etc. Optoacoustic spectra were measured with a resolution of 2 cm^-1^ and an averaging time of 10 000 ms per wavelength between 1700 and 1600 cm^-1^. For comparison and validation, a drop of each solution (5 µL) was measured on an ATR-FTIR spectrometer (ALPHA II from Bruker equipped with a diamond ATR crystal, Bruker Corporation, MA, USA). The FTIR measurements were recorded between 4000 and 400 cm^−1^. Each spectrum was obtained by averaging 150 scans recorded at a resolution of 2 cm^−1^. The spectra were calibrated by measuring a blank sample. The effect of the buffer solution (D_2_O) on the absorption spectra was removed by dividing the sample’s spectra by an independently measured buffer spectrum. The subsequent spectra were then processed using a Savitzky-Golay smooth filter (polynomial order: 5, frame length, smoothing point: 19), baseline correction, and finally, were normalized to 0 – 1. The results are reported in **Fig. 1b-e**, and **Supplementary table 1**.

### Methods section B – HeLa cells

#### Cell culture

A human cervical adenocarcinoma (HeLa) cell line (ATCC: CCL-2) was used to define protein secondary structure composition in living cells. Cells were cultured in Dulbecco’s Modified Eagle Medium (DMEM - Life Technologies, Paisley, GBR) supplemented with 10% fetal bovine serum (FBS, South America, Merck, Darmstadt, DE), 1% L-glutamine and 1% penicillin–streptomycin (Life Technologies, Bleiswijk, NLD).

Cells were grown in a humidified incubator at 37°C, 5% CO_2_, and were passed through trypsin digestion every four days to avoid their reaching full confluence.

#### Spectral imaging analysis

HeLa cells were plated in a custom-made mid-IR dish and cultured until 80% confluence. To interrogate the effect of D_2_O on the absorption spectra and on the image contrast of living HeLa cells in the amide I range, three DMEM media with different % of D_2_O were prepared dissolving DMEM powder (Sigma-Aldrich) in MilliQ water (0% D_2_O), in 70% D_2_O (Sigma-Aldrich) and 30% MilliQ water, and finally in 100% D_2_O. All solutions were supplemented with 3.7 g/L sodium bicarbonate (Sigma-Aldrich), 10% fetal bovine serum, 1% L-glutamine and 1% penicillin–streptomycin. Culture medium was exchanged prior to measurements. HeLa cell micrographs (FOV of 500 μm x 500 μm) were imaged at 1645 cm^-1^, while HeLa cell spectra were acquired between 1700 and 1600 cm^-1^ (step size resolution of 2 cm^-1^ and an averaging time of 10 000 ms per wavelength) in all three media. Absorption spectra were smoothed and normalized with baseline correction (**Fig.1k**) and corresponding second derivative spectra were obtained by interpolation and the Savitsky-Golay method (**Fig.1l**)

#### Contrast-to-Noise Ratio

Contrast-to-noise ratio (***CNR***) is defined as the intensity difference between a point in the sample (***OA_S_***) and a point in the background/cell medium (***OA_ref_***) divided by the peak-to-peak amplitude of the noise level.

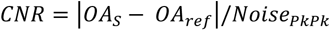

The CNR was calculated in all three HeLa cells micrographs (**Fig. 1g-i**). The noise level is 4.4 mV (pure water), 4.5 mV (70% D_2_O), and 4.7 (100% D_2_O).

#### Contrast profiles of HeLa cells

The plot profiles of HeLa cells in all three media were obtained with ImageJ and normalized as follows:

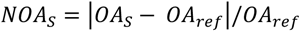

The normalized optoacoustic signal of the cells (***NOA_S_***) was obtained as the intensity difference between the sample optoacoustic signal (***OA_S_***) and the point in the background/cell medium with the lowest contrast (***OA_ref_***) divided by the same point in the background (**Fig.1j**).

#### Cell viability

To interrogate the toxic effect of D_2_O on cell viability, HeLa cells cultured in media composed of different percentages of D_2_O were analyzed with flow cytometry after staining with propidium iodide (PI/RNase, BD Biosciences, San Diego, CA), which easily penetrate dead or damaged cells (see **Supplementary Fig. 3**). Cell viability was determined in three different culture media created by dissolving DMEM powder in 50, 70 and 90% D_2_O and 50, 30 and 10% MilliQ water respectively. DMEM medium created using 100% MilliQ water was used as a control. The media were supplemented with 3.7 g/L sodium bicarbonate, 10% fetal bovine serum, 1% L-glutamine and 1% penicillin–streptomycin as previously described. Cell viability was measured at different time points: 0, 1, 3, 6, 9, 24 hours after exchanging the control (100% MilliQ) medium with a medium partially composed of D_2_O. Cells were then washed with phosphate-buffered saline (PBS) and incubated with PI/RNase (10μg/ml) for 20 minutes in the dark. Cell suspensions were analyzed using Cytoflex LX (Beckman Coulter) flow cytometer and analyzed using FlowJo software.

### Method Section C – MM1.S

#### Cell culture

Human myeloma cells MM1.S (ATCC: CRL-2974) were cultured in RPMI 1640 cell media (Gibco) supplemented with 10% FBS and 1% penicillin-streptomycin. Cells were grown in a humidified incubator at 37°C, 5% CO_2_, and were split 1:5 every four days to avoid cells reaching full confluence.

#### Spectral imaging analysis

MM1.S cells were plated in polylysine (Sigma-Aldrich) coated custom-made mid-IR dishes with a 70-80% confluence. Prior to measurement, the RPMI medium was exchanged with a D_2_O medium obtained by dissolving RPMI-1640 powder (Gibco) in 100% D_2_O. The medium was supplemented with 2g/L sodium bicarbonate (Sigma-Aldrich) and 20% fetal bovine serum. To study the spectral changes caused by misfolded protein formation and consequent apoptosis, we treated myeloma cells for 72 hours with 10µM LEN (Sigma-Aldrich) and with 100 nM BTZ (Velcade, Ratiopharm) (**Figure 2b**). For comparison, myeloma cells not treated with any drug were also plated in the mid-IR dish prior to the measurement (**Fig. 2d**). 10μM LEN solution was prepared by dissolving 250 mg of LEN powder in DMSO (Sigma-Aldrich), while 2.5mg/mL of BTZ (Velcade) was diluted in sterile water to a final concentration of 100nM. After medium exchange, MM1.S cells were imaged at 1650 cm^-1^ at different time points: before addition of any drug (Mt1), after 72 hours of treatment with LEN (Mt2), after 78 hours of LEN treatment and 6 hours of treatment with BTZ (Mt3), and after 96 hours of treatment with LEN and 24 hours of treatment with BTZ (Mt4). At each time point, the amide I spectra of the cells were acquired (1700 – 1600 cm^-1^, step size 2cm^-1^, averaging time 10 000 ms per wavelength). All the spectra acquired were normalized with baseline correction (**Supplementary Fig. 4**). In the differential spectral analysis (**Fig. 2c**), the normalized average spectrum acquired at time 0 (Mt1) was subtracted from the normalized average spectrum acquired at 96 hours (Mt4) (and at the others time points) to detect the appearance of absorption bands assigned to misfolded proteins. For individual drug studies, MM1.S were treated with 25 μM LEN for 144 hours and with 10 μM BTZ for 18 hours.

#### Doxorubicin treatment

In order to confirm the ability of MiROM to selectively detect intermolecular β-sheet structure as a hallmark of misfolded protein formation and consequent apoptosis, MM1.S cells were treated with DOX, a chemotherapy drug that causes an arrest of the cell cycle through DNA intercalation. MM1.S cells were treated with 1μM DOX for 2 hours in 100% D_2_O and spectrally imaged as previously described. MM1.S untreated cells were also spectrally imaged for 2 hours in 100% D_2_O for comparison (**Supplementary Fig. 6**).

#### Computational Analysis

To illustrate MiROM’s sensitivity to protein structure composition, MM1.S spectra were processed with several data analysis tools using the scikit-learn Python package.^58^

##### Principal Component Analysis (PCA)

To estimate the dimensionality of the MiROM spectral dataset, PCA was performed. More than 99.97% of the data variance were explained by the first five principal components.

##### Non-negative matrix factorization (NMF)

Following the dimensionality estimate of the PCA, the spectral data were linearly decomposed into five components via NMF.^59^ The method was chosen in order to obtain physically reasonable features that could be interpreted as partial absorption spectra due to their non-negativity. The NMF was regularized with a combination of Frobenius norm and sparsity promoting L^1^-norm terms. More precisely, the following optimization problem was solved to achieve a decomposition *X* = *WH*, where the rows of the matrix *X* contain the acquired MiROM spectra, the rows of the matrix *H* contain the five NMF components, and the rows of the matrix *W* contain the NMF coefficients of the spectra in *X*:

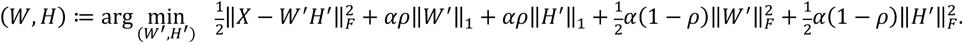

The NMF model was initialized with the non-negative double singular value decomposition method.^60^ A coordinate descent solver was used for the optimization. The tolerance rate and the maximum number of iterations were set to 10^−5^and 2000, respectively. The regularization parameters *α* = 10^−2^ and *ρ* = 10^−3^ were chosen using the L-curve.^61^

##### t-Distributed Stochastic Neighbor Embedding (t-SNE)

To obtain an interpretable visualization of the five-dimensional NMF coefficient data, we performed a non-linear dimensionality reduction with t-SNE to embed the data in two-dimensional space.^62^

## Method Section D – Patient cells

Multiple myeloma cells were biopsied from the bone marrow of twelve patients sensitive or resistant to lenalidomide and bortezomib treatment. Myeloma cells were biopsied according to the guidelines indicated in the ethics vote 672/21 issued by the ethics committee of the Technical University of Munich on 17.11.2021. Informed consent was obtained from all patients prior to measurements. Freshly extracted myeloma cells were plated in polylysine coated mid-IR dishes in RPMI-1640 medium supplemented with 20% FBS, as previously described. In order to investigate the effect of LEN/BTZ treatment on protein secondary structure in patients’ cells, myeloma cells were imaged at 1650 cm^-1^ (FOV 500μm x 500μm, 2μm step size, 50 average) after treatment with 10μM LEN for 72 hours and with 100nM BTZ for 24 hours. At each time point, spectra in the amide I region of 50 cells were acquired (1700 – 1600 cm^-1^, step size 2cm^-1^, averaging time 10 000 ms per wavelength). All spectra acquired were normalized with baseline correction (**Supplementary Fig. 8**) and the differential spectra (**Fig. 3d**) were calculated by subtracting the normalized average spectra acquired at time 0 from the normalized spectra acquired at 72 hours.

### Least – square method

To calculate the treatment response of single cells, we fitted spectra obtained from patients’ cells with the cell medium spectrum by using least – square method. Differential spectra were calculated subtracting the mean value of the fitted spectrum acquired at 0 hours from the spectra at 72 hours. We determined the AUC of the differential spectra in the spectral region between 1638 and 1615 cm^-1^ and compared it with the AUC of untreated cells, using as a threshold the AUC mean value of untreated cells measured at 0 hours.

## Author contributions

F.G. performed all MiROM imaging/spectroscopy experiments, processed the results and prepared the images. M.A.P. designed, built, and characterized the MiROM system. F.G and M.A.P. designed/performed the validation experiments in protein solutions and living HeLa cells. F.G., M.T., M.A.P, F.B. designed the experiment on MM cells and patient samples. F.G. and E.K. performed the FACS experiments. A.N. performed the computational analysis. D.J. supervised the computational analysis. T.Y. helped with the spectral analysis.

A.C. synchronized and automated the imaging system. N.U. helped with the imaging system. F.G. and M.A.P. wrote the manuscript. M.A.P and F.B. supervised the study on MM cells and patient samples, V.N. and M.A.P supervised measurements by optoacoustic detection, M.A.P supervised the whole study. All authors edited the manuscript.

## Acknowledgements

The research leading to these results has received funding from the Deutsche Forschungsgemeinschaft (DFG; German Research Foundation) (project ID 360372040-SFB 1335 and BA 2851/6-1 to F.B.; Gottfried Wilhelm Leibniz Prize 2013; NT 3/10-1 to V.N.), as well as from the European Research Council (ERC) under the European Union’s Horizon 2020 research and innovation programme under grant agreement No 694968 (PREMSOT) to V.N.

The authors thank Dr. Serene Lee, Dr. Elisa Bonnin and Dr. Robert Wilson for assisting with the editing of the manuscript, and Marc Schmidt-Supprian for the use of the Cytoflex LX (Beckman Couldter) flow cytometer.

## Declaration of Interests

V.N. and M.A.P. are founders and equity owners of sThesis GmbH. V.N. is a founder and equity owner of iThera Medical GmbH, Spear UG and I3 Inc. E.K is now an employee of Sandoz, Germany.

## Supplementary figures

**Supplementary Figure 1.**
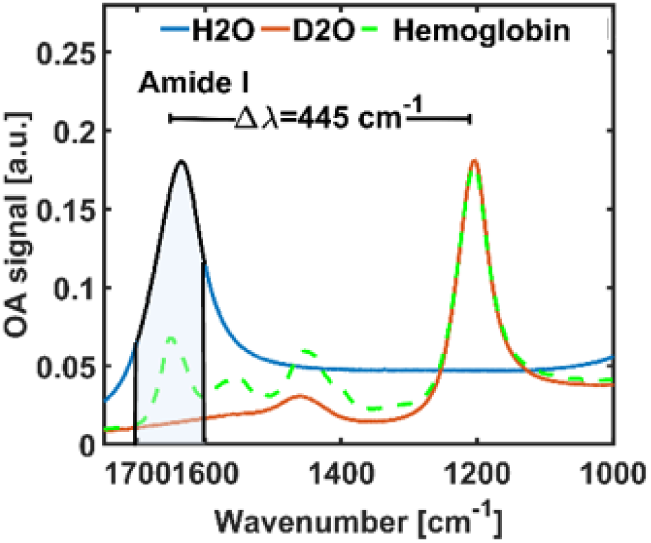
Effect of D2O in the amide I band. Attenuated Total Reflection Fourier Transform InfraRed (ATR-FTIR) spectra of H2O (blue line), D2O (red line) and a D2O albumin solution (green line). The absorption band of H2O overlaps with the amide I band of the hemoglobin solution, preventing its detection. The absorption band of D2O is shifted at 1205 cm^−1^. The use of D2O clarifies amide I absorption region and allows protein detection. The amide I absorption region is highlighted in light blue. OA – Optoacoustic.

**Supplementary Figure 2.**
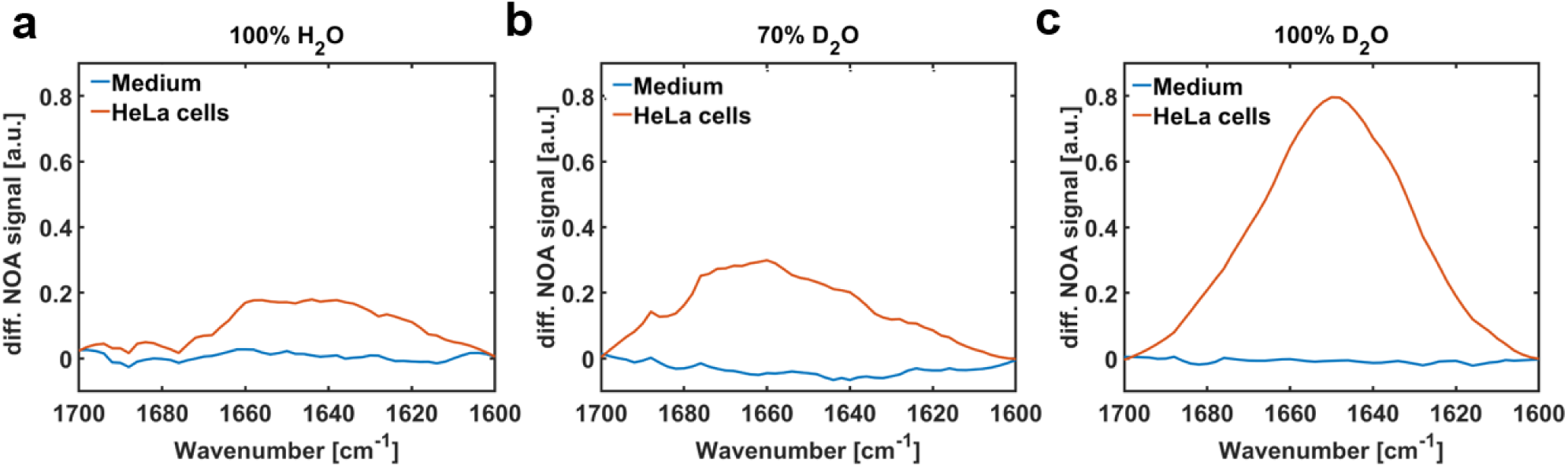
Amide I absorption spectra in HeLa cells. **a)** Amide I absorption spectrum of HeLa cells imaged in a medium composed of 100% H2O. The absorption peak of the cells is two times higher than the baseline (medium). **b)** Amide I absorption spectrum of HeLa cells imaged in 70% D2O. The absorption peak of the cells is three times higher than the baseline (medium). **c)** Amide I absorption spectrum of HeLa cells imaged in 100% D2O. The absorption peak of the cells is eight times higher than the baseline (medium). (n=3 independent experiments). NOA – Normalized Optoacoustic.

**Supplementary Figure 3.**
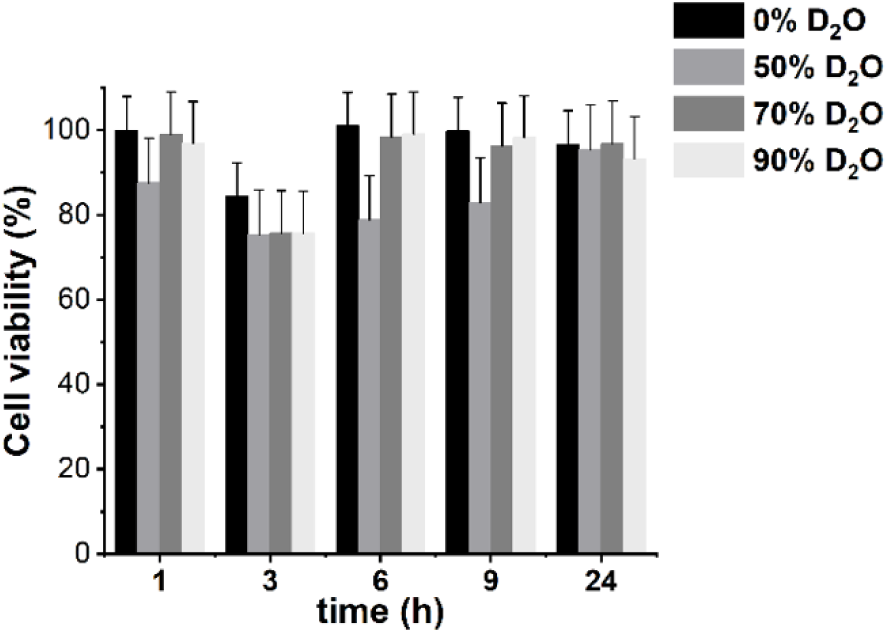
Cell viability. **a)** Propidium iodide (PI) staining for cell viability assessment using Fluorescence-Activated Cell Sorting (FACS) analysis of HeLa cells. HeLa cells were cultured in media containing different percentages of D2O (0, 50, 70 or 90%) and measured at different time points. The use of deuterated water does not appear to change cell viability substantially compared to controls.

**Supplementary Figure 4.**
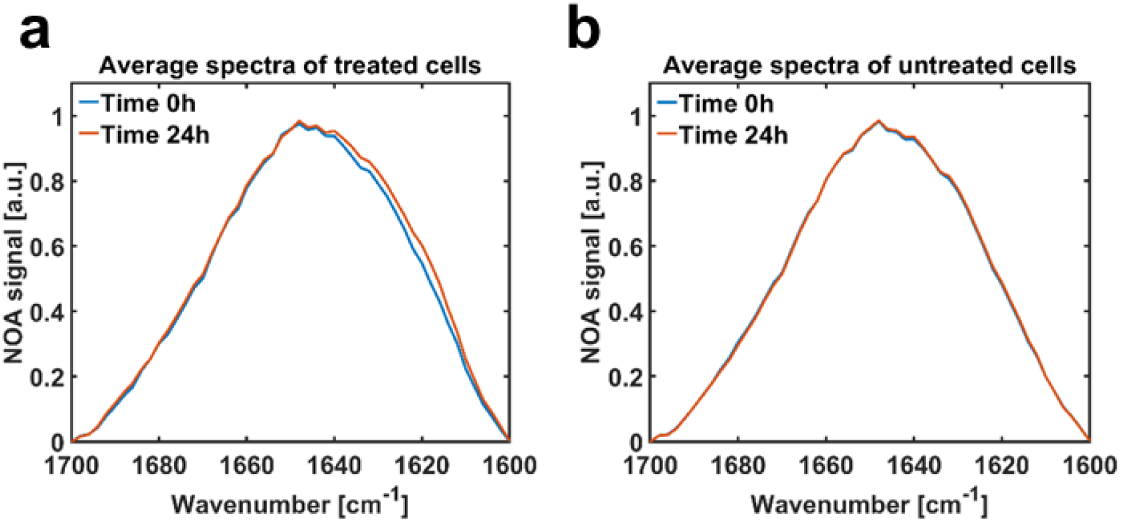
Raw spectra from myeloma cells (MM1.S). **a)** Comparison between the normalized average spectra of 50 myeloma cells at time point 0 (before lenalidomide (LEN) and bortezomib (BTZ) administration) and after 96 hours of LEN/BTZ treatment. **b)** Comparison between the normalized average spectra of 50 untreated myeloma cells at time point 0 and after 96 hours in culture. In (a), the spectrum acquired at 96 hours shows an enlargement towards lower frequencies compared to the spectrum at time 0, while in (b) there is no difference between the average spectra acquired at time 0 and 96 hours after the medium exchange. NOA – Normalized Optoacoustic.

**Supplementary Figure 5.**
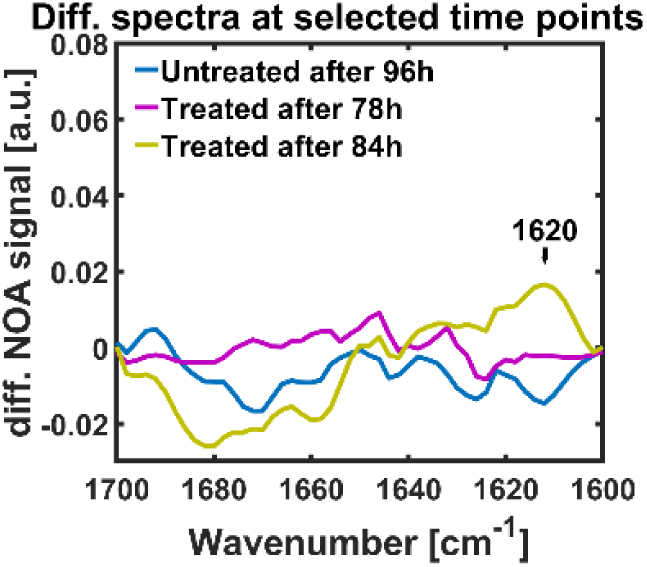
Differential spectra of myeloma cells treated with lenalidomide (LEN) and bortezomib (BTZ). Comparison between the differential spectra of LEN/BTZ treated (in violet and green) and untreated (in blue) myeloma cells acquired in the amide I region at different time points (78h and 84h). The band at 1620 cm^-1^, corresponding to intermolecular β-sheet structures which accumulate in misfolding proteins during the proteasome inhibition treatment, is barely detectable after 84h of treatment. All cells were imaged using a coupling medium composed of 100% deuterated water. NOA – Normalized Optoacoustic.

**Supplementary Figure 6.**
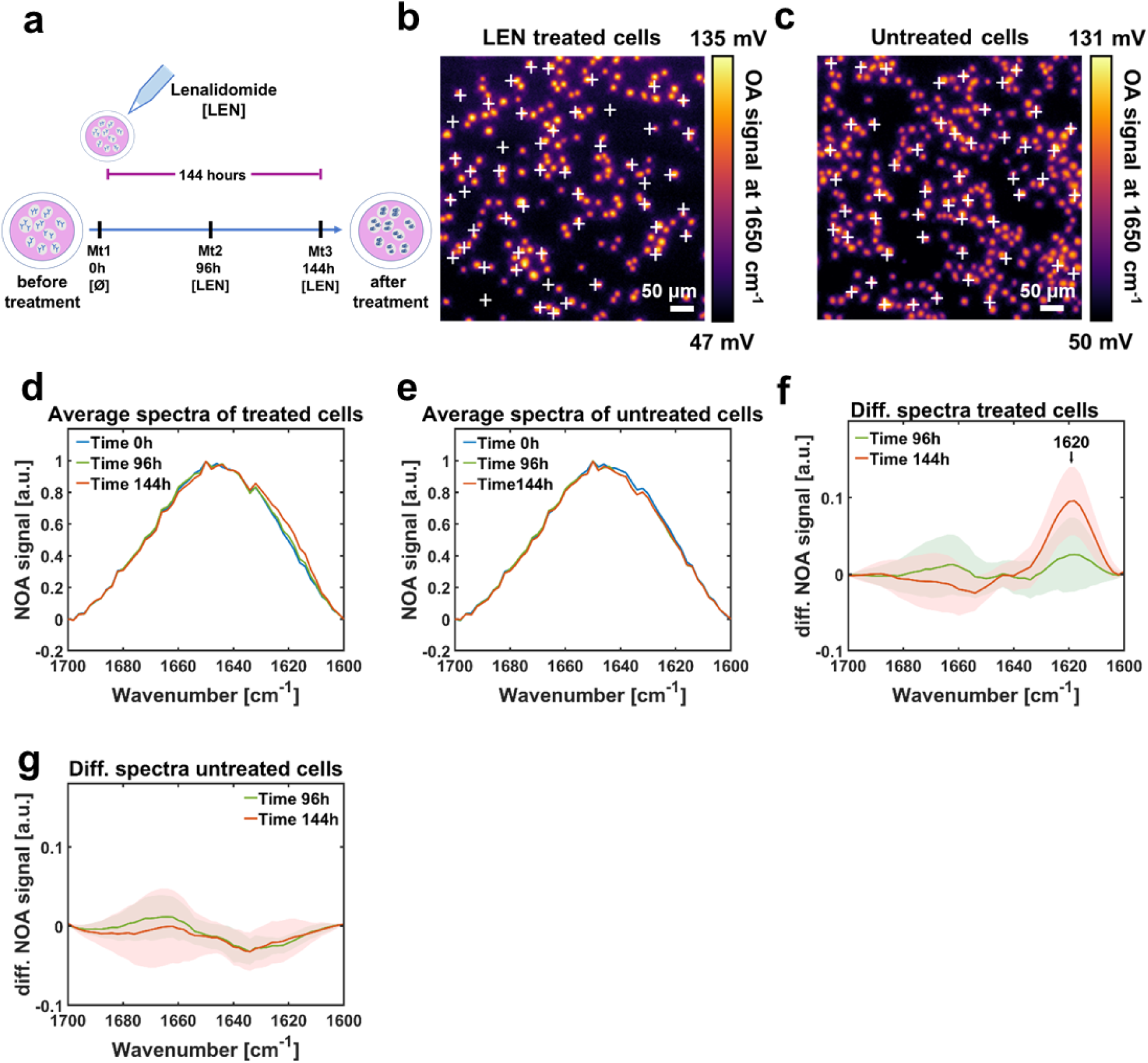
Lenalidomide (LEN) treatment of MM1.S cells. **a)** Schematic diagram of LEN treatment of myeloma cells. **b)** MM1.S cells treated with LEN were imaged after 96 and 144 hours of treatment at 1650 cm^-1^. **c)** For comparison, MM1.S untreated cells were imaged at 1650 cm^-1^ after 96 and 144 hours of being in culture. **d)** Comparison between the normalized average spectra of 50 myeloma cells treated with LEN at time points 0 hours, 96 hours and 144 hours. The average spectrum at 144 hours (in red) shows an enlargement towards lower frequency compared to the spectrum at 0 hours. **e)** Comparison between the normalized average spectra of 50 myeloma untreated cells at time points 0 hours, 96 hours and 144 hours. **f)** Differential spectra of LEN-treated cells at 96 hours (in green) and 144 hours (in blue). Intermolecular β-sheet structures are present in 50% of the cells after 96 hours, (green band at 1620 cm^-1^), and in 100% of the cells after 144 hours (red band at 1620 cm^-1^). **g)** Differential spectra of untreated cells at 96 and 144 hours. The intermolecular β-sheet band at 1620 cm^-1^ is absent in the spectra of untreated cells (n=3 independent experiments). OA – Optoacoustic. NOA – Normalized Optoacoustic.

**Supplementary Figure 7.**
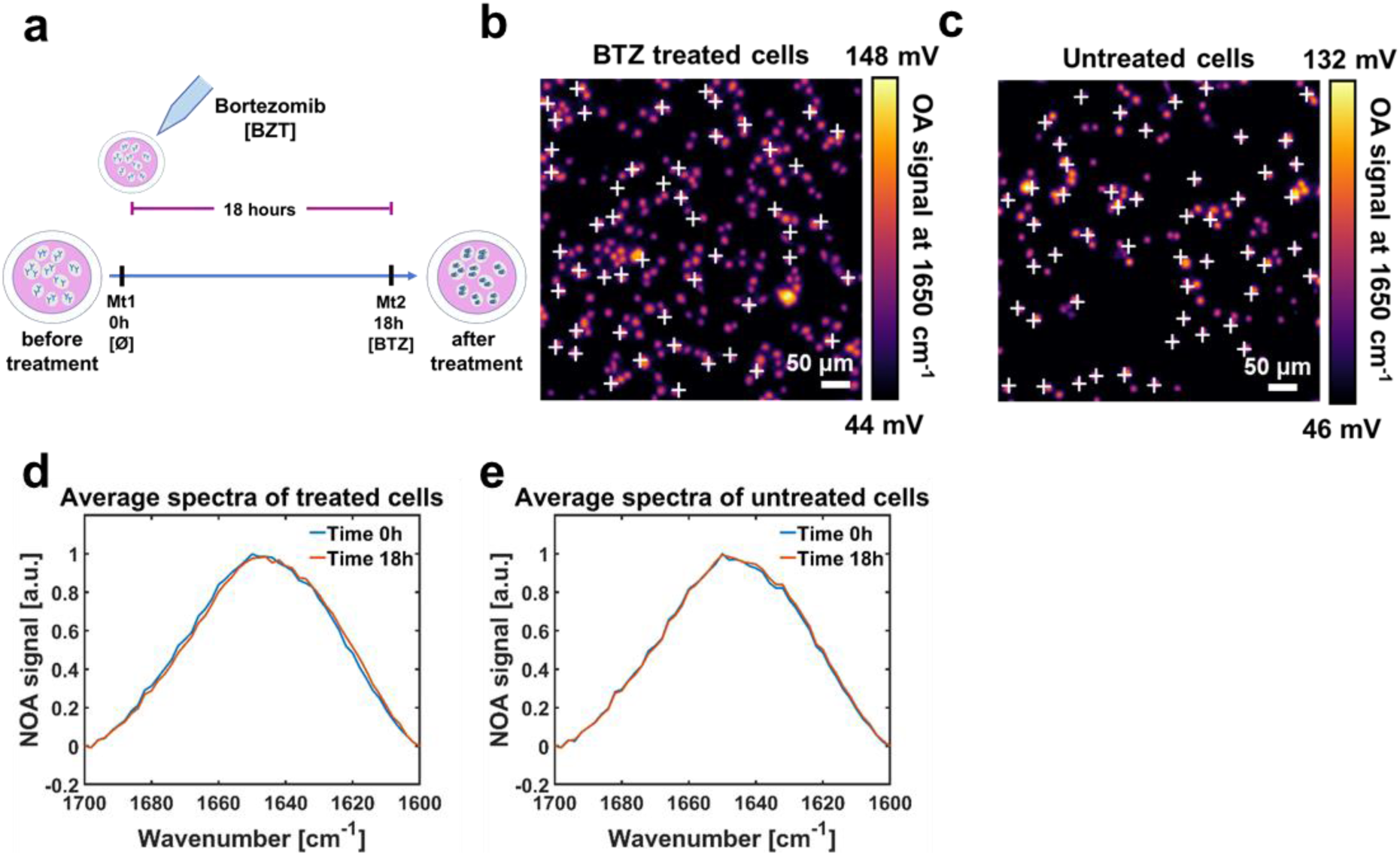
Bortezomib (BTZ) treatment of MM1.S cells. **a)** Schematic diagram of BTZ treatment of myeloma cells. **b)** MM1.S cells treated with BTZ were imaged at 1650 cm^-1^after 18 hours of treatment. **c)** For comparison, MM1.S untreated cells were imaged for 18 hours at 1650 cm^-1^. **d)** Comparison between the normalized average spectra of 50 myeloma cells treated with BTZ at time point 0 hours and at 18 hours. The average spectrum acquired after 18 hours (in red) shows a small enlargement towards lower frequency compared to the spectrum acquired at 0 hours. **e)** Comparison between the normalized average spectra of 50 myeloma untreated cells at 0 and 18 hours. (n=3 independent experiments). OA – Optoacoustic. NOA – Normalized Optoacoustic.

**Supplementary Figure 8.**
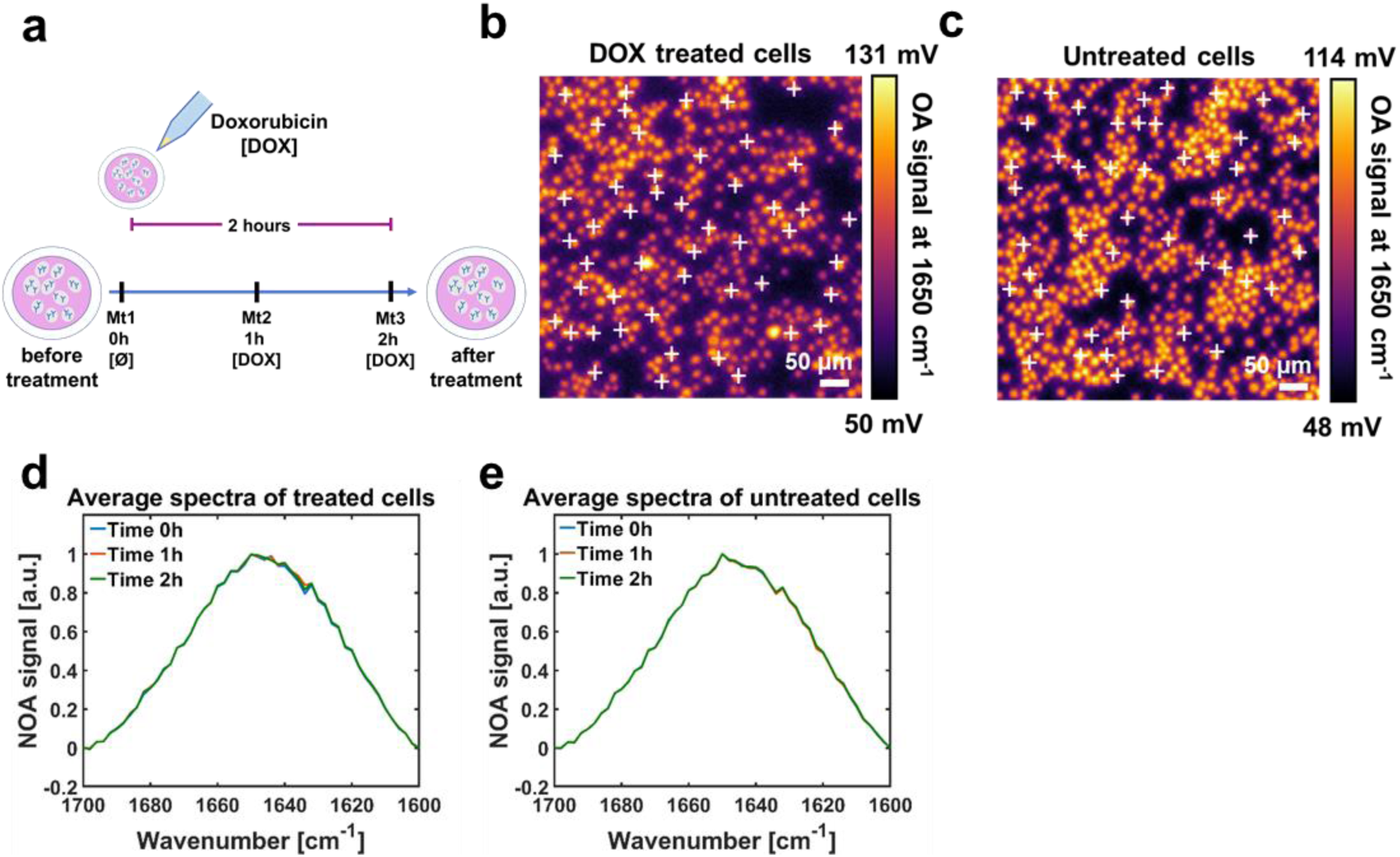
Doxorubicin (DOX) treatment of MM1.S cells. **a)** Schematic diagram of DOX treatment of myeloma cells. **b)** MM1.S cells treated with DOX were imaged for 2 hours at 1650 cm^-1^. **c)** For comparison, MM1.S untreated cells were imaged for 2 hours at 1650 cm^-1^. **d)** Comparison between the normalized average spectra of 50 myeloma cells treated with doxorubicin at time points 0 hours, 1 hours and 2 hours. **e)** Comparison between the normalized average spectra of 50 myeloma untreated cells at time points 0 hours, 1 hours and 2 hours. Spectra in (d) and (e) are comparable. (n=3 independent experiments). OA – Optoacoustic. NOA – Normalized Optoacoustic.

**Supplementary Figure 9.**
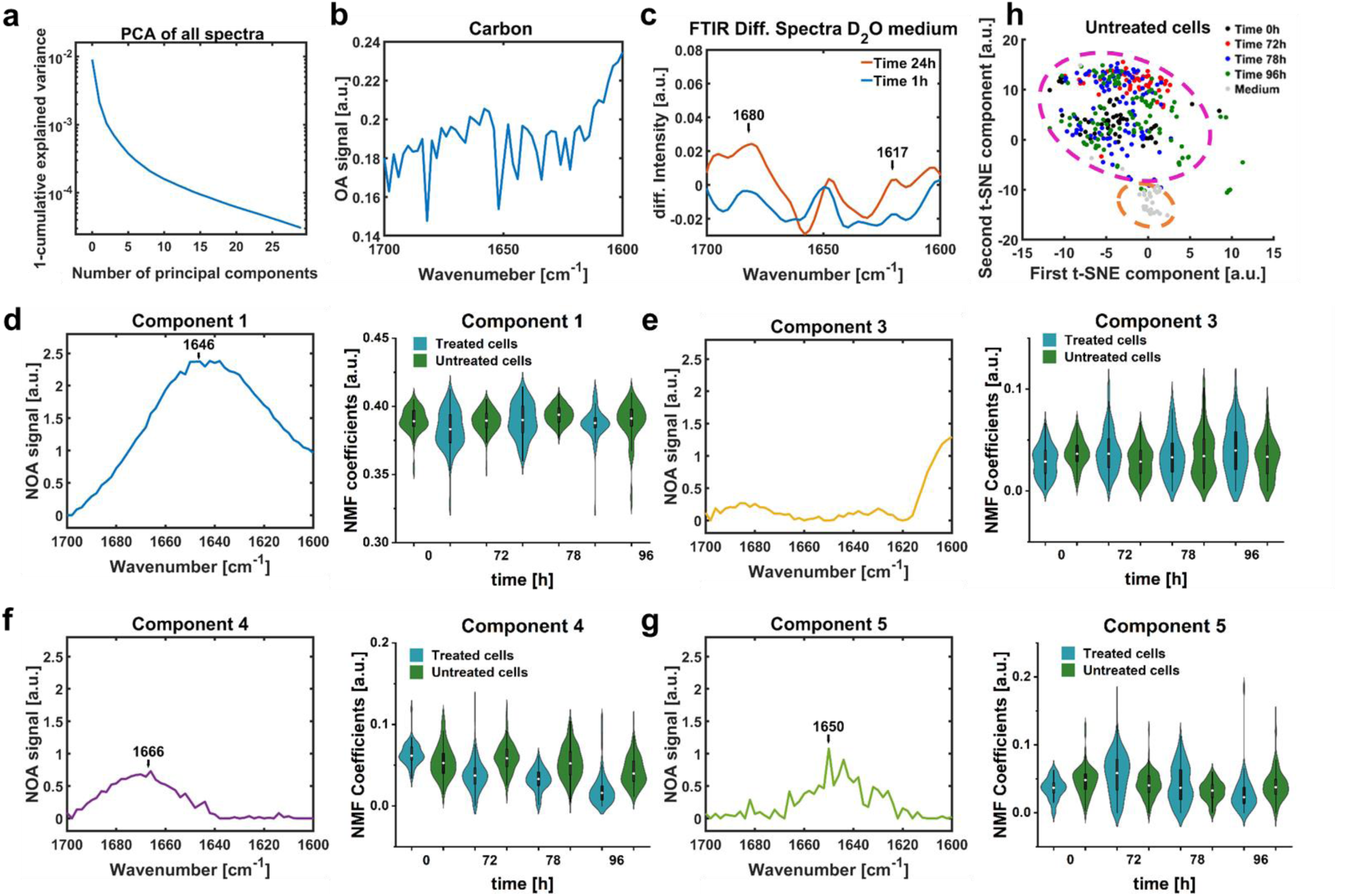
Principal Component Analysis (PCA) and Non-negative Matrix Factorization (NMF) components. **a)** PCA of all spectra obtained from myeloma cells (MM1.S) treated with lenalidomide (LEN) and bortezomib (BTZ) and untreated MM1.S cells. **b)** Emission profile of the laser. **c)** Fourier Transform InfraRed (FTIR) spectra show a band around 1680 cm^-1^ characteristic of H/D exchange. **d-g)** NMF components and violin plots from kernel density estimate the time evolution of the components’ coefficients for LEN/BTZ treated and untreated myeloma cells. **h)** A t-Stochastic Neighbor Embedding (t-SNE) map representing the distribution of all 5 components in untreated myeloma cells. (n=700 spectra). NOA – Normalized Optoacoustic.

**Supplementary Figure 10.**
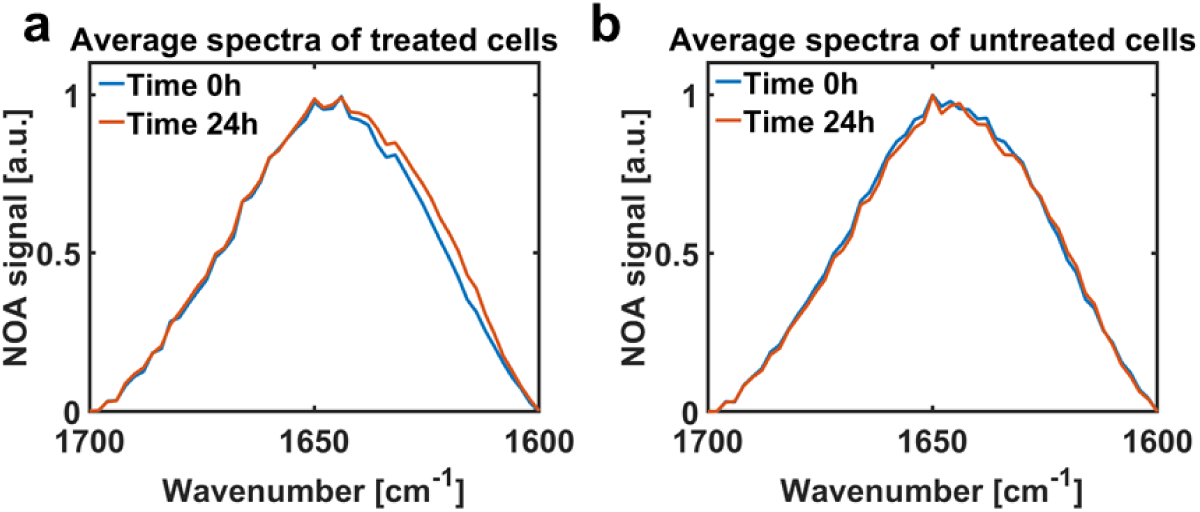
Spectra of myeloma cells from patients sensitive to lenalidomide (LEN) and bortezomib (BTZ). **a)** Comparison between the normalized average spectra of myeloma cells from 50 patients at time point 0 (in blue, before LEN and BTZ administration), and after 72 hours of LEN treatment and 24 h of BTZ treatment (in red). **b)** Comparison between the normalized average spectra of 50 untreated myeloma cells purified from a patient at 0 hours (in blue), and 72 hours (in red) after the medium exchange. In (a), the spectrum acquired at 72 hours shows an enlargement towards lower frequencies compared to spectrum at 0 hours, while in (b) there is no difference between the average spectra acquired at 0 hours after 72 hours in culture. (n=10 patients). NOA – Normalized Optoacoustic.

**Supplementary figure 11.**
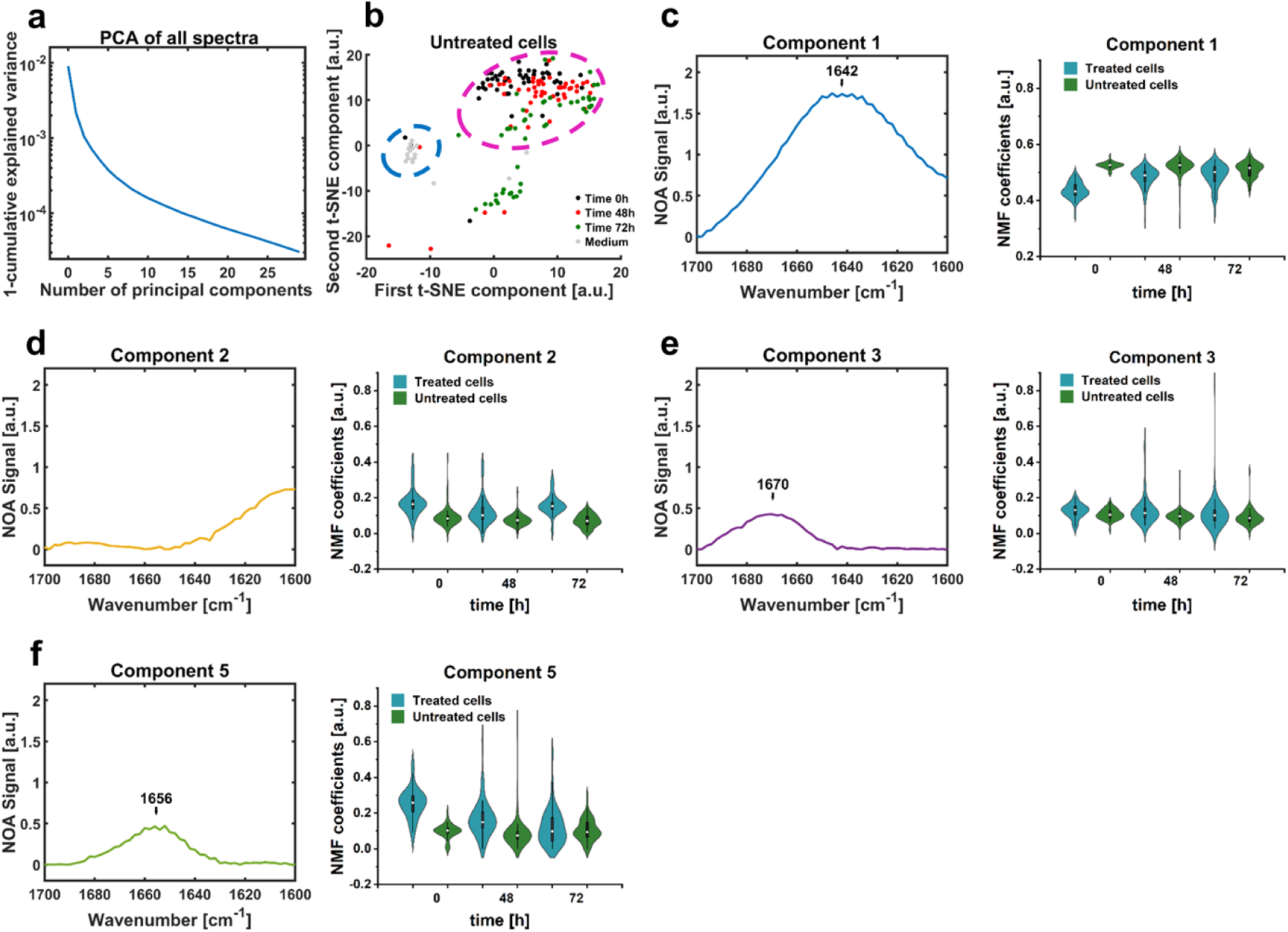
Image monitoring and computational analysis of untreated myeloma cells purified from a patient. **a)** Principal Component Analysis (PCA) performed on myeloma cells treated with lenalidomide (LEN) and bortezomib (BTZ) and untreated myeloma cells. **b)** t-Stochastic Neighbor Embedding (t-SNE) map representing the distribution of the 5 components identified by Non-Negative Matrix Factorization (NMF) in untreated cells. **c-f)** NMF components and violin plots from kernel density estimate the time evolution of the components’ coefficients for LEN/BTZ treated and untreated myeloma cells. Component 1 at 1642 cm^-1^ (c) refers to the medium, component 2 (d) describes the emission profile of the laser, component 3 at 1670 cm^-1^ (e) and component 5 at 1656 cm^-1^ (f) are associated with α-helix structures. (n=400 spectra). NOA – Normalized Optoacoustic.

**Supplementary figure 12.**
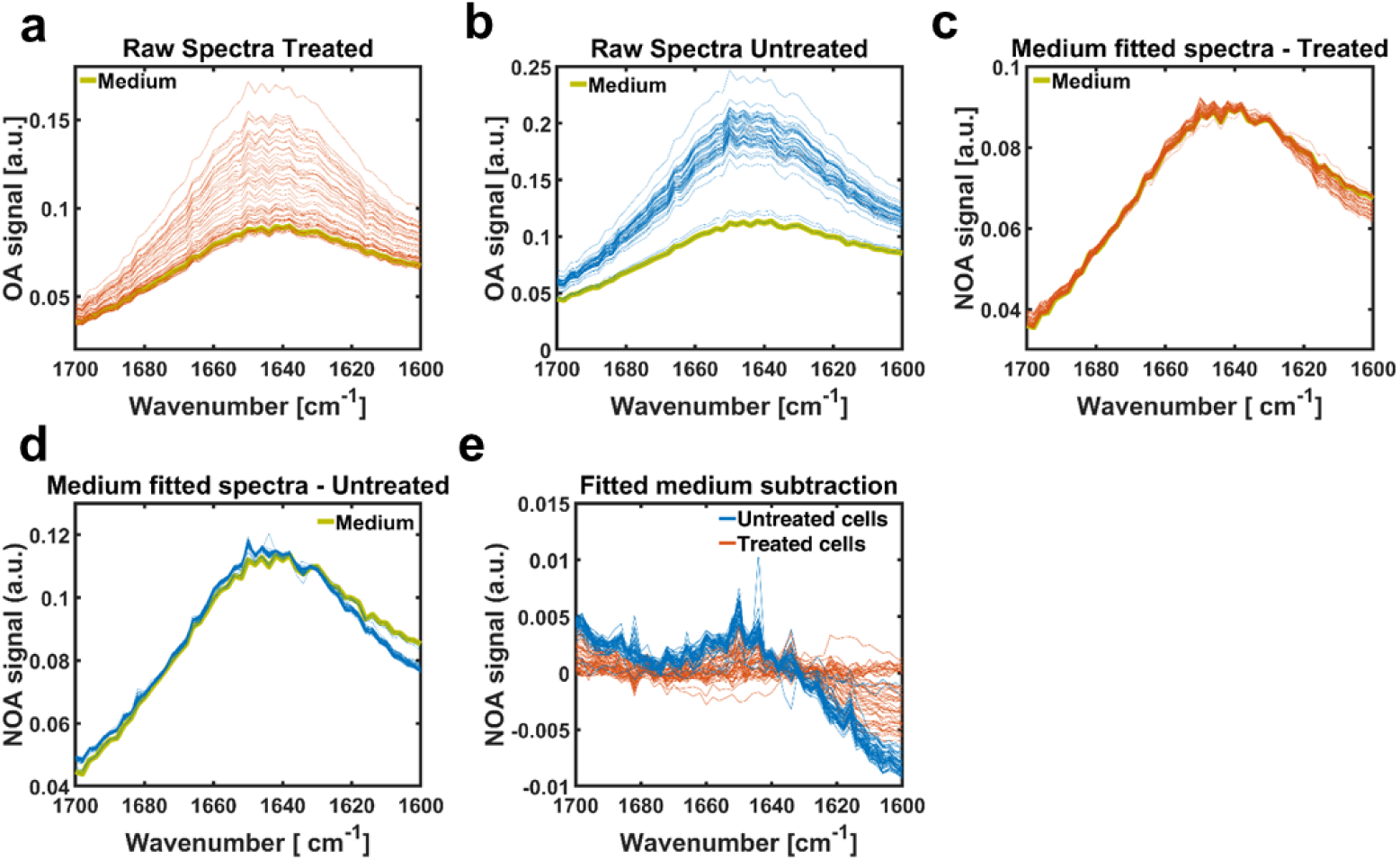
Least – square method. Raw spectra of lenalidomide (LEN) and bortezomib (BTZ) treated **(a)** and untreated **(b)** patient cells with the corresponding medium spectrum (in green). Spectra of LEN/BTZ treated **(c)** and untreated **(d)** patient cells fitted with their corresponding medium spectrum using the curve least squares fitting method. **e)** Spectra of LEN/BTZ treated and untreated cells after subtraction of the fitted medium spectrum. **f)** Comparison between differential spectra of LEN/BTZ treated and untreated single cells extracted from a patient sensitive to LEN and BTZ. LEN/BTZ treated cells show an increase of signal in the region of the intermolecular β-sheet structure (1638 – 1615 cm^-1^). OA – Optoacoustic. NOA – Normalized Optoacoustic.

**Supplementary figure 13.**
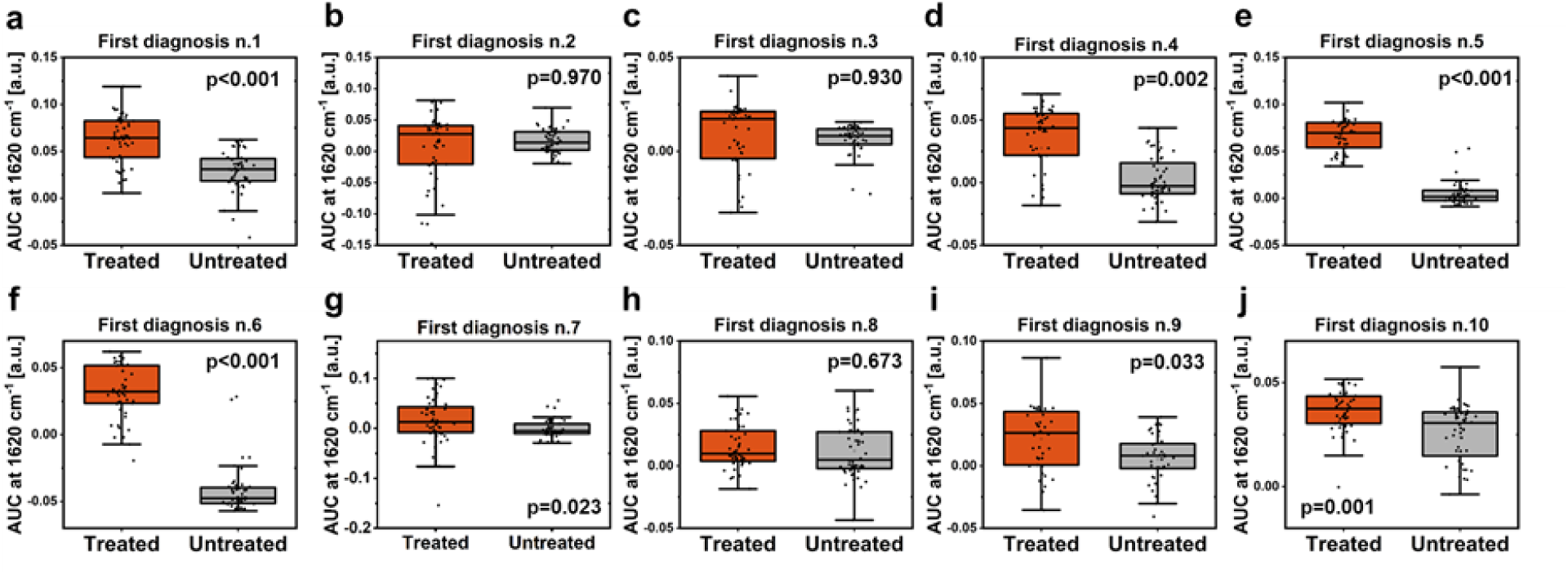
Single-cells response of lenalidomide (LEN) and bortezomib (BTZ) sensitive myeloma patients. **a-j)** Boxplot representing the Area Under the Curve (AUC) of the band at 1638 – 1615 cm^-1^ of the amide I differential spectra obtained in treated and untreated cells extracted from the bone marrow of 10 LEN and BTZ sensitive myeloma patients. The corresponding percentage responses are reported in **Table 3**. P values from paired sample *t*-test.

**Supplementary figure 14.**
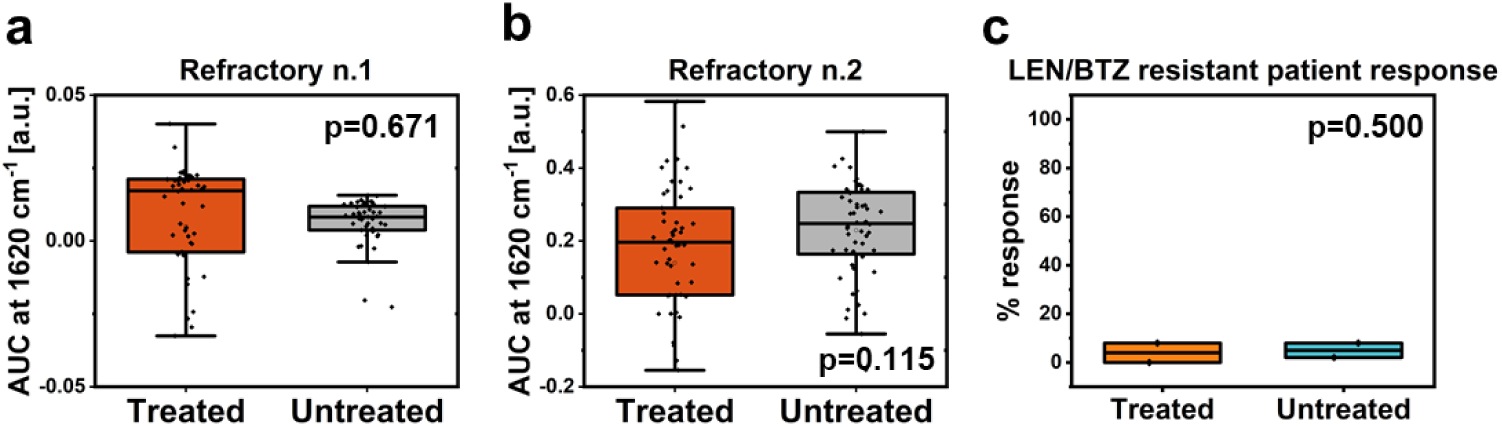
Single-cell response of lenalidomide (LEN) and bortezomib (BTZ) resistant myeloma patients. **a, b)** Boxplot representing the Area Under the Curve (AUC) of the band at 1638 – 1615 cm^-1^ of the amide I differential spectra obtained in LEN/BTZ treated and untreated cells extracted from the bone marrow of 2 refractory myeloma patients. The corresponding percentage responses are reported in **Table 3**. **c)** Boxplots representing the percentage response (%) of LEN/BTZ treated and untreated cells analyzed from 2 independent patients resistant to LEN and BTZ. P values from paired sample *t*-test.

